# Dynamic variability in apoptotic threshold as a strategy for combating fractional killing

**DOI:** 10.1101/375915

**Authors:** Baohua Qiu, Jiajun Zhang, Tianshou Zhou

**Affiliations:** Key Laboratory of Computational Mathematics, Guangdong Province; School of Mathematics, Sun Yat-Sen University, Guangzhou, 510275, P. R. China

## Abstract

Fractional killing, which is a significant impediment to successful chemotherapy, is observed even in a population of genetically identical cancer cells exposed to apoptosis-inducing agents. This phenomenon arises not from genetic mutation but from cell-to-cell variation in the activation timing and level of the proteins that regulate apoptosis. To understand the mechanism behind the phenomenon, we formulate complex fractional killing processes as a first-passage time (FPT) problem with a stochastically fluctuating boundary. Analytical calculations are performed for the FPT distribution in a toy model of stochastic p53 gene expression, where the cancer cell is killed only when the p53 expression level crosses an activity apoptotic threshold. Counterintuitively, we find that threshold fluctuations can effectively enhance cellular killing by significantly decreasing the mean time that the p53 protein reaches the threshold level for the first time. Moreover, faster fluctuations lead to the killing of more cells. These qualitative results imply that dynamic variability in threshold is an unneglectable stochastic source, and can be taken as a strategy for combating fractional killing of cancer cells.

## Introduction

Resistance to chemotherapeutic agents remains a major obstacle to effective cancer treatment. Much effort has been devoted to understanding mechanisms of resistance to improve the therapeutic effect. Previous studies considered that drug resistance emerges due to specific mutations in a subset of tumor cells, and it is those mutated cells that survive chemotherapy treatment (1). However, recent experimental investigations into genetically identical populations of tumor cells exposed to apoptosis-inducing agents revealed that drug resistance also emerges through mechanisms of non-genetic mutations, often through stochastic fluctuations in key factors in response to drugs. Drug resistance means that some cells are killed while others survive during treatment. This phenomenon is known as fractional killing (2).

Single-molecule measurement technologies have shed much light on the underlying molecular mechanisms of cell-to-cell variability in fractional killing (2–5). For example, experiments verified that genetic mutations in BCR-ABL can give rise to fractional killing of cancer cells or lead to the drug ineffectiveness, but in two-thirds of cases no genetic mutations was found (6). In the case of apoptosis mediated by tumor necrosis factor-related apoptosis-inducing ligand, it is common that a part of tumor cells of a clonal population are killed while the others survive (2). In human cell lines, fractional killing arises from cell-to-cell variability in the timing and probability of death, and this variability is thought to originate from the differences in the levels of the proteins that regulate receptor-mediated apoptosis (3). Another experimental observation is that the cell-to-cell variability in p53 dynamics can result in fractional killing, where a cell’s death probability depends on the time and level of p53 and the cell must reach a dynamically fluctuating threshold to execute apoptosis (referring to Fig. 1*A*)(3). In spite of these case-to-case experimental efforts, how non-genetic variability in the timing and level of key proteins regulating apoptosis impacts fractional killing of cancer cells remains to be not fully understood, and model efforts are required to address this intriguing yet important issue, especially in the case of dynamic variability in apoptotic thresholds.

**FIGURE 1.**
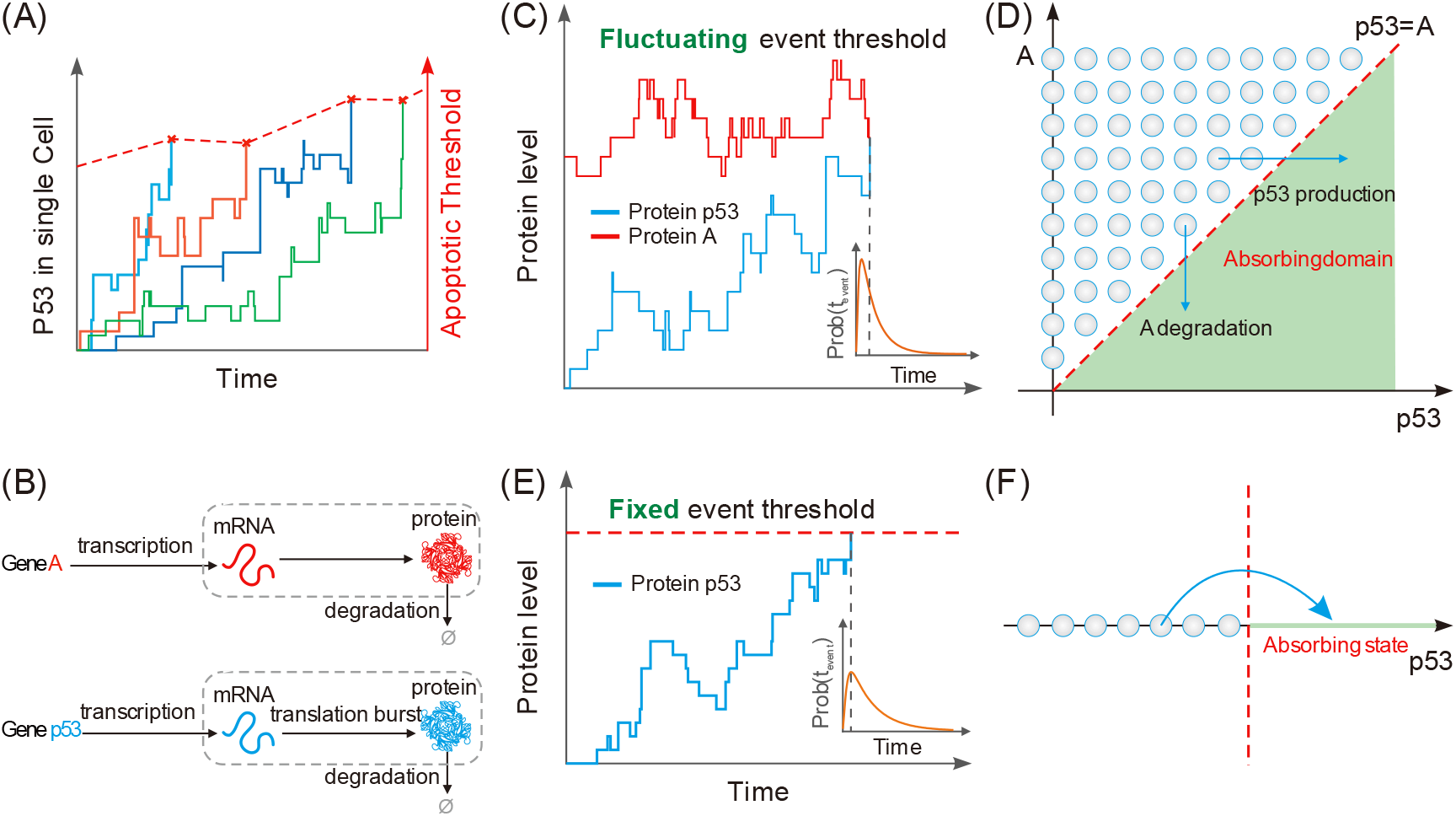
Threshold crossing can be modeled as a FPT problem. (A) A realistic example involving FPT, where cells must reach a threshold level of *p*53 to execute apoptosis and this threshold increases with time (3). (B) Two gene models, where the model for gene p53 assumes that the gene is expressed in a burst manner whereas that for gene A assumes that the gene is expressed in a constitutive manner. (C) A FPT problem with a dynamically fluctuating threshold, where *A*(*t*) represents the threshold curve that *p*53(*t*) hits for the first time, and the inset shows the FPT distribution. (D) A one-dimensional FPT problem with a dynamically fluctuating threshold is transformed into a two-dimensional FPT problem with a fixed threshold. The shadow region represents an absorbing domain of FPT in the (*p*53, *A*) plane, defined by 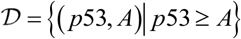, where an event is triggered once p53 crosses *A*, the line of *A* = *p*53 represents the boundary of region 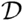, and arrows represent the possible directions of threshold crossing. (E) The expression level of gene *p*53 changes over time, where the dashed red line represents a critical threshold that the gene product crosses, and the inset shows the FPT distribution. (F) Schematic for an absorbing domain of FPT, where the empty circles with arrow represents threshold crossing.

As is well known, for many cancer types, the p53 transcription factor is a key regulator in the cellular response to DNA damage induced by chemotherapy (7). Experimental evidence supports that increasing upstream p53 abundance can trigger the transcription of multiple genes in various downstream programs including cell apoptosis and cell-cycle arrest (8). Previous works suggested a threshold mechanism where the choice between different programs depends on p53 protein levels (9,10). In the corresponding models, low levels of p53 triggers cell-cycle arrest and high levels of p53 leads to apoptosis. Subsequently, some studies (11,12) showed that the dynamics of p53 plays a role in the specificity of the response with pulsed p53 favoring DNA repair and cell-cycle arrest genes and sustained p53 triggering activation of senescence and apoptotic genes. Recently, Paek, *et al.*(3), used live cell imaging to investigate the role of p53 dynamics in fractional killing of colon cancer cells in response to chemotherapy. They showed that both surviving and dying cells reach similar levels of p53, implying that cell death is not determined by a fixed p53 threshold. Conversely, a cell’s probability of death depends on the time and levels of p53. They also showed that cells must reach a threshold level of p53 to execute apoptosis and this threshold increases with time. The increase in p53 apoptotic threshold is due to drug-dependent induction of anti-apoptotic genes, predominantly in the inhibitors of apoptosis family. These quantitative experiments call for a corresponding modeling effect that addresses the question of how dynamic variability in apoptotic threshold affects fractional killing of cancer cells.

In order to address this issue, we first formulate complex fractional killing processes as a FPT problem and then analyze a simplified model of stochastic p53 dynamics, where the cancer cell is killed only when the p53 expression level crosses a dynamically fluctuating apoptotic threshold. Analytical calculations are performed for the FPT distribution in this model. Counterintuitively, we find that dynamic variability in the apoptotic threshold can effectively enhance cellular killing by significantly decreasing the mean time that the p53 protein reaches the threshold level for the first time. In addition, we also find that faster fluctuations can lead to the killing of more cells. These qualitative results indicate that stochastic fluctuations in apoptotic thresholds are an unneglectable noisy source that can facilitate killing of cancer cells. Therefore, tuning this variability would be a potential strategy for combating fractional killing and thus improving drug efficacy.

## Materials and methods

### Modeling fractional killing processes as a FPT problem

Fractional killing generally results from the crosstalk between complex apoptosis pathway and complex survival pathways. These complexly structured and heterogeneous processes as well as the paucity of experimental data hamper effort to construct detail models. However, fractional killing processes are essentially threshold-crossing events. For clarity and in order to reveal the essential mechanism of how dynamically fluctuating thresholds impact the dynamics of threshold crossing, we consider a toy model of gene regulation (referring to Fig. 1*B*), where a timing event is triggered once the expression level of a gene (its product is denoted by p53) crosses the expression level of another gene (its product is denoted by A) for the first time. Indeed, simple mathematical models are important tools toward to understanding the essential mechanisms of important biological processes such as cell apoptosis and interpreting experimental phenomena (13–16). They can also provide guidelines for experimental designs with a growing interest in combining clinical and molecular data.

Specifically, we use a stochastic model of p53 gene expression to investigate the effect of stochastic fluctuations in apoptotic thresholds on fractional killing of cancer cells. This model explicitly takes into account “molecular noise” in the p53 protein that regulates apoptosis in the emergence of drug resistance during treatment, and dynamic variability in the apoptotic threshold. Indeed, dynamically fluctuating thresholds have a strong biological background and are ubiquitous in biological regulatory systems. For example, consider a representative activity function of Hill type (17–19) Activation = *Z^n^*/(*Z^n^* + *K^n^*) where *n* is a Hill coefficient, and *K* = *k*_−_/*k*_+_ represents a threshold (in fact a dissociation constant) of variable *Z*. In general, reaction rate *k*_+_ or *k*_−_ is regulated by external signals that are stochastically generated due to biochemical reactions, e.g., *k*_−_ = *α*_1_*S* + *α*_0_ where *S*, an external signal, is stochastically generated, *α*_0_ and *α* are positive constants. In this case, *K* is a dynamically fluctuating threshold of *Z*. For a gene whose expression has to reach a dynamically fluctuating threshold level, expression noise and threshold fluctuations can all lead to variability in the event timing. This raises questions: how these two stochastic origins impact threshold crossing, and which regulatory strategies can control variability in the event timing. Most previous studies have focused on the first-passage properties of stationary threshold crossing (17,18,20–22), while comparatively very few studies have investigated how a dynamically fluctuating threshold impacts timing precision and arrival time theoretically (22–26) or experimentally (3).

Now, we formulate the stochastic temporal timing of events as a problem of FPT to a dynamically fluctuating threshold. Denoting by *p*53 and *A* the p53 protein and the apoptotic threshold, respectively. Cells must reach an apoptotic threshold level of p53 to execute apoptosis, and this threshold fluctuates dynamically with time. Let {*p*53 (*t*)}_*t*≥0_ be a temporally homogeneous stochastic process with initial *p*53_0_, and {*A* (*t*)}_*t*≥0_ represent a dynamically fluctuating threshold (boundary or barrier) with initial *A*_0_. Without loss of generality, we set *p*53_0_ < *A*_0_. This setting is natural since *A* represents the critical threshold that *p*53 will cross. Note that the union of two trajectories *p*53(*t*) and *A*(*t*), (*p*53(*t*), *A*(*t*)) constitutes a new system. Define *T* as the time that trajectory *p*53 (*t*) hits trajectory *A*(*t*) for the first time, that is,

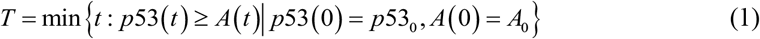

which is called the first passage time (FPT) (20–22). Apparently, *T* is a random variable since both *p*53(*t*) and *A*(*t*) are stochastic, referring to Fig. 1*A*. The left issue is how the distribution of *T* including statistical quantities is correlated to stochastic dynamics of *p*53(*t*) and *A*(*t*). Our basic idea is to transform a one-dimensional FPT problem with dynamically fluctuating threshold into a two-dimensional FPT problem with a fixed boundary.

For analysis convenience, we consider a two-gene expression model to mimic *p*53 induced tumor cell apoptosis. Assume that the *p*53 molecules are produced in a burst manner whereas the *A* molecules are generated in a constitutive manner. We use the produced counts of protein molecule *A* to construct a stochastically fluctuating threshold to the molecular number of protein *p*53. Let *p*53(*t*) ∈ {0,1,2, …} denote the level of protein *p*53 at time *t*, and assume that protein *p*53 is generated with a Poisson rate 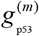 (where superscript (*m*) means that feedback regulation is considered, but it may be omitted in the absence of feedback regulation) and degrades at a constant rate *d*_p53_. The translation burst approximation is based on the assumption of short-lived mRNAs, that is, each mRNA degrades instantaneously after producing a burst of *B* protein molecules, where *B* follows a geometric distribution (27–30), that is

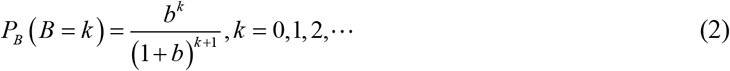

where *b* represents the mean protein burst size. Then, we can show *P_B_*(*B* ≥ *k*) = (*b*/(1 + *b*))*^k^*, *k* = 0,1,2, ⋯. In what follows, we denote *P_B=k_* ≡ *P_B_* (*B* = *k*) and *P_B≥k_* ≡ *P_B_*(*B*≥*k*) for simplicity. Similarly, *A*(*t*) ∈ {0,1,2,…1} represents the level of protein *A* at time *t*, and is assumed to follow a Poisson distribution with two characteristic parameters 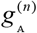 (where the meaning of superscript (*n*) is similar to that of superscript (*m*)) and *d_A_* representing respectively the transcription and degradation rates of protein A when *A*(*t*) = *n*. The time evolution rule of (*p*53(*t*), *A*(*t*)) is defined as follows: (*p*53(*t*), *A*(*t*)) starting from (*p*53(*t*) = *m*, *A*(*t*) = *n*)) with *m* < *n* at time *t* is updated through the following probabilities of timing events in the infinitesimal time interval (*t*, *t* + *dt*]

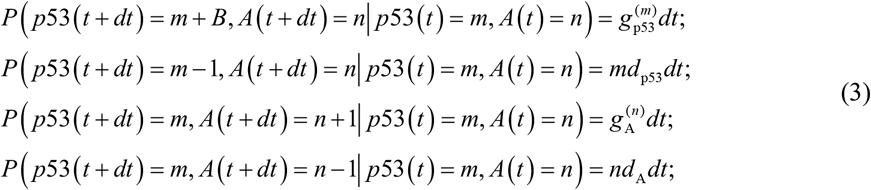

The apoptosis event occurs if the cumulative number of protein *p*53 molecules exceeds the number of protein A molecules. Noted that threshold *A*(*t*) is not a constant but fluctuates over time. In what follows, we will investigate the effect of the noise in *A*(*t*) on threshold-crossing events. In addition, we will compare the FPT characters between cases of fluctuating (i.e., *A*(*t*) stochastically changes) and fixed (i.e., *A*(*t*) = constant) thresholds. Note that the more threshold-crossing events are, the fewer cancer cells are killed and the more cancer cells are killed otherwise.

### Stochastic model formulation for FPT problems to a fluctuating threshold

The problems of FPTs to fluctuating thresholds arises in many scientific fields such as biology, statistics and engineering. However, in contrast to fixed threshold FPT problems, it seems to us that there have been no methods to handle fluctuating threshold FPT problems. For the above example, we successfully transform a one-dimensional FPT problem with dynamically fluctuating threshold into a two dimensional FPT problem with a fixed boundary. This transform can easily be extended to more complex cases. It is worth pointing out that this transform can easily be extended to a more complex case.

Specifically, we introduce an absorption domain 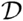, which consists of those points (*p*53, *A*) that satisfy *p*53 ≥ *A*. that is, 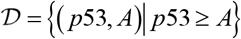. Let *P_m,n_* represent the probability that a two-dimensional system is at state (*m, n*) at time t, that is,

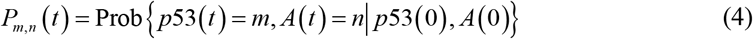

For convenience, *P_m,n_*(*t*) is sometimes denoted by *P_S_* (*t*), i.e., *P_S_* (*t*) = *P_m,n_*(*t*), where *S* = (*p*53, *A*) represents state. Note that the survival probability is equal to the sum of the probabilities of all the states that do not belong to the absorbing region, i.e., 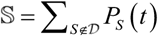. Denote by *f_T_*(*t*) the probability density function for the FPT, that is, *f_T_*(*t*) = Prob {*T* ≤ *t*}.

The relation between the protein molecules *p*53 and *A* can be considered as a trajectory in the domain {(*p*53, *A*)| *p*53(*t*) < *A*(*t*)}. The corresponding forward master equation (FME) describing the time evolution of protein pair *p*53 and *A* can be described as the following master equation (20,29,30)

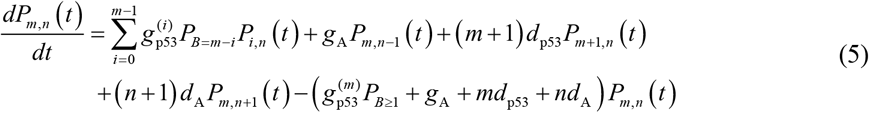

where *m* < *n*. Then, the FPT distribution can be formally expressed as (22,31,32)

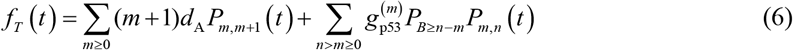

In numerical simulation, we constrain *n* = 1,2, ⋯, *C* (where *C* is a pre-given positive integer), implying that *m* = 0, 1, 2, ⋯, *C* − 1.

### Statistical quantities of FPT distribution

Although the FPT distribution in principle provides complete characterization of the threshold crossing timing, we are particularly interested in the lower-order statistical moments of FPT distribution (*f_T_*(*t*)). Starting from a general FME, we can obtain analytical formulas for the first and second-order moments of FPT. For this, we first establish the relation between distribution *f_T_*(*t*) and state *S*, and then give the formal expression of *f_T_*(*t*). Assume that all states {*S*(*t*)}_*t*≥0_ with *S*(*t*) = (*p*53 (*t*), *A*(*t*)) constitute a Markov process. Then, we have a FME of the form (20,29,30)

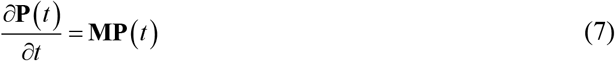

where **P**(*t*) is a column vector consisting of all *P_s_*(*t*), and **M** is a certain linear operator, depending a process of interest. Note that every component of **P**(*t*) is the probability that the system {*S*(*t*)}_*t*≥0_ arrives at the absorbing domain 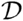 at time *t*, and that **M** is actually a state transition matrix. Let *P*(*S_f_, t*|*S, t*_0_) be the probability that the state *S*(*t*) reaches the absorbing state *S_f_* at time *t*, given the initial state *S* = *S*(*t*_0_) at time *t*_0_ with *S*(*t*_0_) = (*p*53_0_, *A*_0_) (*t*_0_ = 0 can be set). And let 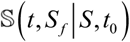 be the survival probability that the trajectory {*S*(*t*)~ starting from *S* at time *t*_0_ has not yet been absorbed to state *S* at time *t*, that is, 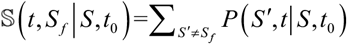. By definition, we have 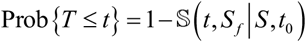. Thus, the probability density function of the FPT is given by (See the S1 Supporting Information for more details.)

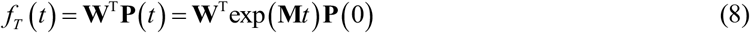

where **W** is the column vector of the transition rates from all accessible states to the absorbing state and the superscript T represents transpose (22,31,32).

Once the probability density function of the FPT, *f_T_*(*t*), is given or found, we can show that raw moments of random variable *T* are given by (see the S1 Supporting Information for derivation)

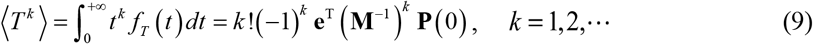

where **e**^T^ = [1, 1,⋯,l] is a constant vector. This indicates that the moments of FPT can be calculated directly based on the FME once the initial transition probabilities **P**(0) are set. In particular, the *timing mean*, i.e., the mean FPT (MFPT) is calculated according to

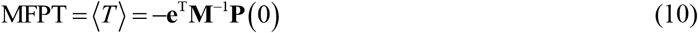

which is a statistical quantify of our main interest. In addition, the intensity of the noise in *T* (defined as the ratio of variance over the square of mean), which represents the *timing variability* or reflects the precision in the event timing, is calculated according to

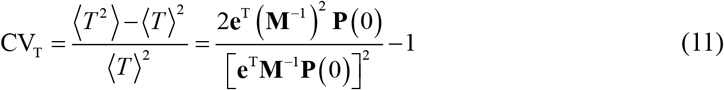

And other higher order moments such as skewness and kurtosis can also be formally given, detailed in the S1 Supporting Information. From Eq. (8) to Eq. (11), we know that to calculate statistical quantities, the key is to calculate the inverse of matrix **M**. Note that the more the 〈*T*〉 is, the fewer are the cancer cells killed, and the smaller the CV_T_ is, the more precise is the threshold crossing.

It should be pointed out that the number of protein *p*53 or *A* molecules may be infinite in theory, implying that **M** in Eq. 7 is an infinite-dimensional matrix. Therefore, Eq. 9, Eq. 10 and Eq. 11 have only theoretical significance since they give only the formal expressions of FPT distribution and statistical quantities. Owing to such infinity, the FPT problem we study here is essentially different from a traditional FPT problem in which matrix **M** is finitely dimensional due to the fixed threshold. The infinite-dimensional FPT problem is in general intractable, and it is thus needed to develop computational methods. Here we propose a so-called truncation approach to solve this tough problem. This approach is developed based on the finite state projection (33), seeing the next subsection for details.

### An efficient method for solving the problem of FPT to a fluctuating threshold

#### (A) Algorithm description

For clarity, we use the above FPT model to introduce our truncation method, which can be generic and applied to more complex cases.

First, the finite state projection approach (33) tells us that matrix **M** can be replaced by a *k×k* sub-matrix **M**_*k*_, so that the approximation 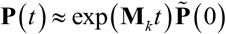 holds, where 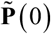 replaces the original **P**(0) in some order. As a result, the state vector (*p*53_*i*_, *A_i_*) constitutes a finite state projection, where *i* ∈ {1,2,⋯,*k*}.

Second, we define 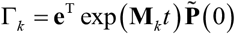. which represents the sum of the components of vector **P**(*t*). According to the finite state projection approach, we can prove that if Γ_*k*_≥1 − *ε* with *ε* being a small positive number, we have

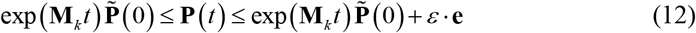

Based on the above analysis, we develop the following truncation algorithm:

**Inputs** Propensity functions and stoichiometry for all reactions.

Initial probability density vector 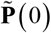
Final time of interest, *t_f_*.
Total amount of acceptable error, *ε*.
Initial finite set of states, (*p*53_0_, *A*_0_).
Initialize a counter, *k* = 0.

**Step 0** Calculate **M**_*k*_ = Submatrix (**M**), which depends on (*p*53_*k*_, *A_k_*), and

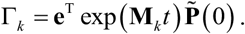

**Step 1** If Γ_*k*_≥ 1 − *ε*, **Stop**.

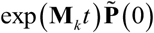 approximates the probability **P**(*p*53_*k*_,*A_k_,t_f_*) with error *ε*.

**Step 2** Add more states, (*p*53_*k*+1_, *A*_*k*+1_) = expand ((*p*53_*k*_, *A_k_*)), and take *k* ← *k*+1. Increment *k* and return to **Step 1**.

#### (B) The expression of matrix M

Owing to the effectiveness of the truncation approach proposed above (referring to Fig. 2), we may assume that matrix **M** is finitely dimensional (otherwise, we use finitely dimensional matrix, **M**_*k*_). To derive the expression of matrix **M** in the above gene model, we consider a special absorbing domain

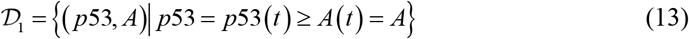

**FIGURE 2.**
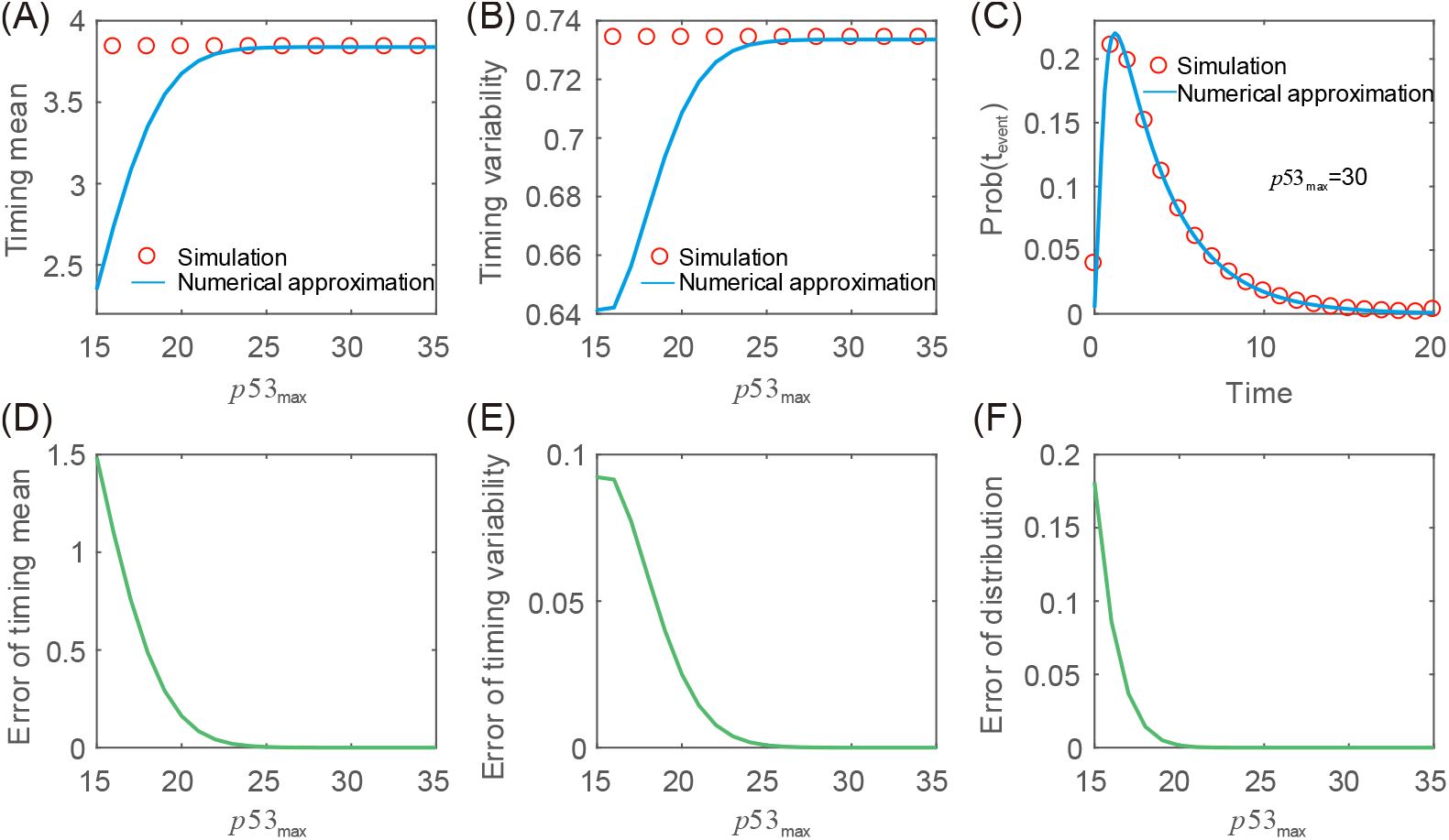
Verification of the effectiveness of the truncation algorithm with the model described in Fig 1(B). (A) Timing mean as a function of *p*53_max_, (B) Timing variability as a function of *p*53_max_. (C) The FPT distribution for *p*53_max_= 30. (D) The difference between simulation and approximate results of timing mean as a function of *p*53_max_. (E) The difference between exact and approximate results of timing variability as a function of *p*53_max_. (F) The Kullback-Leibler divergence between exact and approximate FPT distribution as a function of *p*53_max_. In (A)-(F), empty circles represent the results obtained by the Gillespie stochastic simulation and are therefore viewed as “exact”, whereas the curve represents the result obtained by our algorithm and are therefore viewed as “approximate”. The parameter values are *g*_p53_ = 5, *d*_p53_ = 1, *b*= 1, *g*_A_ = 10, *d*_A_ = 1, the mean value of *A* (i.e., the mean threshold) is set as *A*_threshold_ = 10, and the range of *p*53 is set as *p*53_max_ = 15 ~ 35.

The S1 Supporting Information performs analysis for other three kinds of absorbing domains.

Introduce the numerical cutoffs for the numbers of proteins *p*53 and *A* respectively: *p*53_max_ for *p*53(*t*) and *A*_max_ for *A*(*t*), and assume *p*53_max_=*A*_max_=*C*(a known integer) without loss of generality. Therefore, *m* ∈ {0,1,2,3,…,*C*−1} and *n* ∈{1,2,3,…,*C*} for the above gene model (see Fig. 1*F*). That means that the corresponding finite state-space for birth-death process can be considered as follows

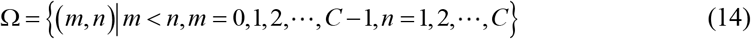

For convenience, we rewrite vector **P**(*t*) as **P** = [**P**_*,1_,**P**_*,2_,**P**_*,3_,⋯,**P**_*,*C*_]^*T*^, where **P**_*,*k*_ represents **P**_*,*k*_= [*P*_0,*k*_, *P*_1,*k*_,⋯,*P*_*k*−1,*k*_,0,⋯,0], *k* ∈ {1,2,⋯,*C*}, and time *t* is omitted. Also for convenience, we introduce an operator, denoted by 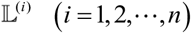, which acts on matrix with the operation rule being as follows: 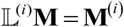. where **M**^(*i*)^ is a matrix whose order is the same as that of **M** but some components are possibly zero, e.g., if **M** = (*a_ij_*)_3×3_, we can have 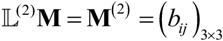, where (*b_ij_*)_2×2_ = (*a_ij_*)_2×2_ and the other elements are equal to zero. Thus, matrix **M** above can be expressed as the following form

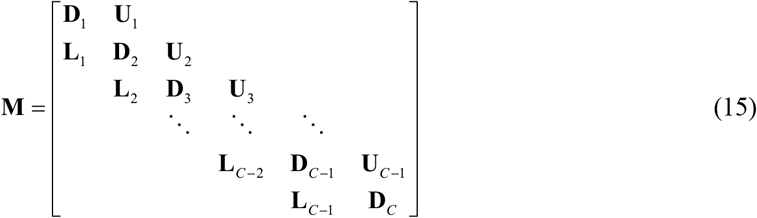

where

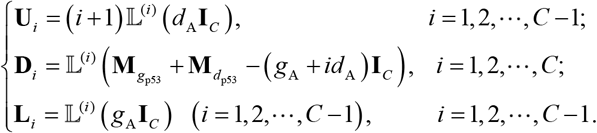

Here **I**_*c*_ is an identity matrix; matrix **M**_*d*_p53__=*d*_p53_diag([1,2,…,*C*−1],1)−*d*_p53_diag([0,1…,*C*−1],0), where symbol diag (**v**, *k*) represents that the elements of vector **v** are placed on the *k*th diagonal. Note that *k*=0 corresponds to the main diagonal, *k*>0 corresponds to above the main diagonal, and *k*<0 corresponds to below the main diagonal; matrix **M**_gp53_ = *g*_p53_**M**_burst_, where 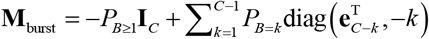. In the presence of feedback, implying that *g*_p53_ depends on the molecule number (*m*) of protein *p*53, we have 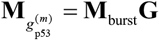, where 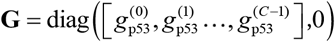 (See the S1 Supporting Information for the formal expressions of these matrices).

#### (C) The expression of FPT distribution *f_T_*(*t*)

Given a numerical cutoff (*C*), the FPT distribution in Eq.8 can be rewritten as (see the S1 Supporting Information for more details)

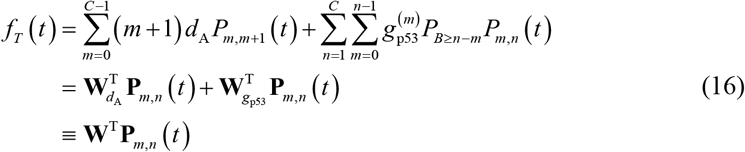

where 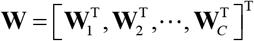 with 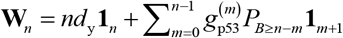, *n*=1,2⋯,*C*. Here we define a column vector of length *C*, **1**_*i*_= (0,⋯ 0,1,0,⋯, 0)^T^ in which the only *i*th element is equal to 1 and other elements are all zero.

For a given **P**(0), the mean FPT 〈*T*〉 and the noise CV_*T*_ can be calculated by Eq.10 and Eq.11 respectively, where a key step is to calculate the inverse of matrix **M** through Eq.15. In contrast, the FPT distribution *f_T_*(*t*) is easily calculated through Eq.16. In a word, through the calculation of these quantities, we can analyze characteristics of timing events with fluctuating thresholds, including the first passage time and variability in the timing. For a sake of simplicity, we will not consider feedback regulation implying that 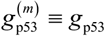 and 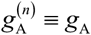 are independent of *m* and *n*.

#### (D) The effectiveness of the truncation algorithm

In order to verify the effectiveness of the truncation algorithm proposed above, we perform numerical calculation using the gene model described in Fig. 1 *B*. Numerical results are shown in Fig. 2.

Figs. 2*A* and 2*B* demonstrate the dependence of the mean FPT and the timing variability on the cutoff of the protein *p*53 number, respectively. We observe with increasing the cutoff constant, both can well approximate the “exact” values (empty circles) obtained by the Gillespie stochastic algorithm (34) with increasing the cutoff constant. For example, the approximate mean FPT is nearly equal to the exact mean FPT at *p*53_max_≈ 23 whereas the approximate timing variability is nearly equal to the timing variability at *p*53_max_≈ 25. Fig. 2*C* demonstrates the distribution of FPT for a particular cutoff constant *p*53_max_= 30.

Figure 2*D* demonstrates the dependence of the difference between the exact mean FPT obtained by the Gillespie stochastic algorithm (34) and the approximate mean FPT obtained by the finite state projection on the cutoff constant *p*53_max_. We observe that this difference quickly tends to zero as the cutoff constant is beyond some value. The similar change tendency holds for timing variability, referring to Fig. 2*E*. In addition, Fig. 2*F* more clearly demonstrates that two kinds of FPT distributions are in agreement since the Kullback–Leibler divergence (35) between them tends to zero as the cutoff constant *p*53_max_ increases, further verifying the effectiveness of the proposed truncation algorithm.

In the remainder of this article, we will use the numerical method above to compute two statistical quantities of FPT distribution, the *timing mean* and the *timing variability*. The former characterizes the response time of *p*53reaches the apoptosis threshold, i.e., the shorter the *timing mean*, the more cells are killed. The latter quantifies the precision in threshold crossing timing. Lower variability implies more robust cell-killing strategy.

## Results

### Stochastic fluctuations in apoptotic thresholds can accelerate tumor cell apoptosis

First, we introduce the conception of event threshold for convenience. By event threshold we mean that the average level of protein *A* is equal to the ratio of the generation rate (*g*_A_) over the degradation rate (*d*_A_), e.g., the event threshold *A*_threshold_=*g*_A_/*d*_A_. Thus, an event threshold is nothing but a fixed threshold in the deterministic case (i.e., in the case of no fluctuations). In the case of fluctuations, however, an event threshold may not be equal to the fixed threshold due to the effect of stochastic fluctuations.

Then, in order to show how fluctuating thresholds impact the timing of events, we plot Fig. 3 where numerical results in the case of fixed threshold are also shown for comparison. From Fig. 3 *A*, we observe that the curve for the dependence of mean FPT on event threshold in the case of fluctuations is always below the curve for the dependence of the mean FPT on event threshold in the case of no fluctuations, implying that threshold fluctuations always shorten the time that the regulatory proteins reach a critical threshold, or accelerate the response of intracellular events to external cues. As a supplement of Fig. 3*A*, Fig. 3*B* demonstrates the dependence of the difference between the mean FPT in the case of fixed threshold and that in the case of fluctuating threshold on event threshold. We observe that this difference is a monotonically increasing function of event threshold. The insets show two special FPT distributions, which correspond respectively to one empty circle and one triangle indicated in the figure.

**FIGURE 3.**
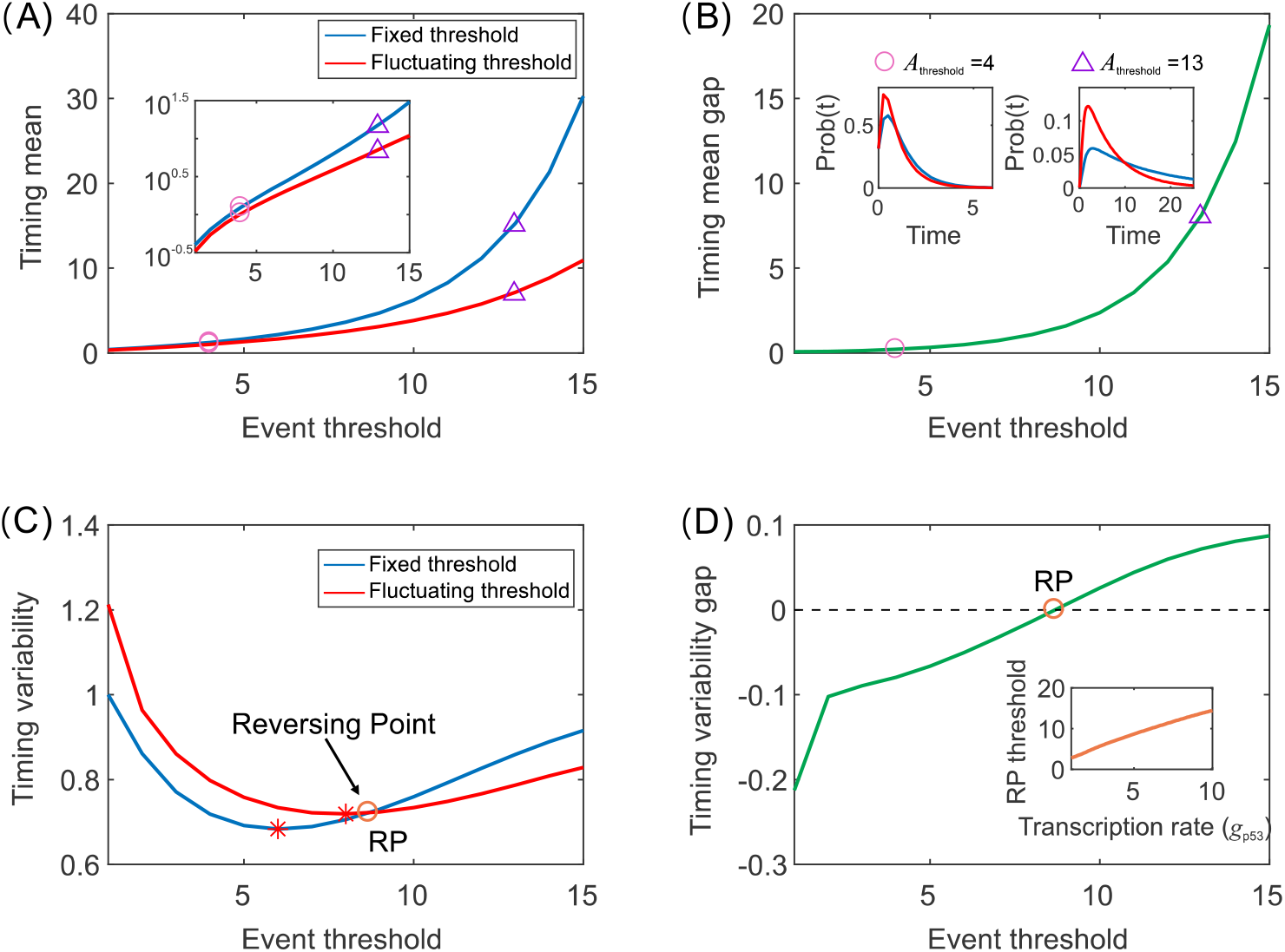
Comparison between the effects of fixed and fluctuating thresholds on timing. (A) Timing mean as a function of event threshold for two different kinds of thresholds, where the inset shows timing mean as a function of event threshold on the logarithmic scale. (B) A different demonstration of the results in (A), showing the difference of timing mean in the case of fixed threshold minus that in the case of fluctuating threshold, where insets show FPT distributions for two different event thresholds (indicated by empty circle and triangle) corresponding to *A*_threshold_= 4 and *A*_threshold_= 13. (C) Timing variability as a function of event threshold for two different kinds of thresholds, where the empty circle is the crossing point of two curves, and stars represent the critical threshold that makes timing variability reach the minimum. (D) As a supplement of (C), the difference of timing variability in the case of fixed threshold minus that in the case of fluctuating threshold, where the inset shows the critical threshold as a function of the transcription rate. In (A) and (C), the parameter values are set as *g*_p53_= 5, *d*_p53_= 1, *b*= 1, *p*53_max_ = 30, the fixed threshold is *A*_threshold_ = 10, and the fluctuating threshold corresponds to *g*_A_= 10 and *d*_A_= 1. The inset in (D) corresponds to *d*_A_= 1, *b* = 1, *g*_A_=10, *d*_A_= 1, *g*_p53_, =1 ~ 10, and *p*53_max_=30.

Figure 3*C* demonstrates how event threshold impacts variability in the timing. From this figure, we observe that there is a critical event threshold (denoted by RP) such that the timing variability in the case of fluctuating threshold is smaller than that in the case of fixed threshold, as the event threshold is beyond RP but the former is larger than the latter as the event threshold is below RP. In other words, for a high event threshold, threshold fluctuations can reduce the timing variability or can raise the precision in the timing. In addition, we observe that there is an even threshold such that the variability in the timing is least in both cases of threshold (referring to the star indicated). This implies event threshold can make the timing precision reach optimality in both cases of fixed and fluctuating thresholds. Fig. 3*D* is a different demonstration of the results in Fig. 3*C*, showing that the difference between the timing variability in the case of fixed threshold and that in the case of fluctuating threshold is a monotonically increasing function of event threshold. The inset shows the dependence of the critical event threshold on the transcription rate *g*_p53_, demonstrating that the critical event threshold increases with *g*_p53_.

In short, Figs. 3*A* and 3*C* show our main results, that is, dynamic fluctuations in threshold can accelerate the response of intracellular events to external cues by shortening the time that the regulatory proteins reach the critical threshold for the first time, fluctuations in high event thresholds can raise the precision in timing by reducing timing variability, and there is an even threshold such that the timing variability reaches optimality in both cases of fixed and fluctuating thresholds. These results imply that threshold fluctuations are an important factor affecting the timing of events, and also that dynamic variability in apoptotic threshold can facilitate the killing of cancer cells.

### Fast fluctuations in apoptotic thresholds can lead to killing of more cancer cells

In the above subsection, we have seen that fluctuations in thresholds have important influences on event timing. However, factors leading to such fluctuations may be diverse. Here we focus on investigating the effects of timescales on timing mean and timing variability.

First, we give the definition of timescale. In our model, if the production rate (*g*_p53_ or *g*_A_) and the degradation rate (*d*_p53_ or *d*_A_) of protein *p*53 or *A* are simultaneously enlarged by *α*_p53_ or *α*_A_ times, then the factor *α_p53_* or *α*_A_ is defined as the timescale of protein *p53* or *A*. In general, the larger the factor *α*_p53_ or *α*_A_ is, the larger are the fluctuations in protein *p*53 or *A*. Therefore, *α*_p53_ or *α*_A_ is an important factor leading to fluctuations in protein *p53* or *A*. Small *α_p53_* or *α*_A_ corresponds to slow fluctuations whereas large *α*_p53_ or *α*_A_ to fast fluctuations. We can prove that the variability in event timing depends only on the ratio of *α*_A_ over *α*_p53_, independent of their sizes. See theS1 Supporting Information for details.

Then, we investigate the influence of timescales on mean FPT and variability in the even timing. Numerical results are shown in Fig. 4. Specifically, Figs. 4*A* and 4*D* demonstrate how timescales of proteins *p*53 and *A* together affect the mean FPT and timing variability, respectively. We observe that the mean FPT in the case of small *α*_p53_ is always larger than that in the case of large *α*_p53_, independent of *α*_A_ (referring to the red dashed line). For a fixed yet small *α*_p53_ (referring to the black dashed line), the mean FPT in the case of small *α*_A_ is also larger than that in the case of large *α*_A_. These imply that two kinds of timescales (internal for *p53* and external for *A*) can all significantly impact the mean FPT. Moreover, a larger external timescale leads to a less mean FPT for small internal timescale, but a smaller internal timescale leads to a larger mean FPT for small external timescale. On the other hand, this relationship is different in the case of timing variability, referring to Fig. 4D. We observe that there is a strip region (indicated by orange) in the plane of *α*_p53_ and *α*_A_, such that the variability in timing is largest. More precisely, if two boundary lines of this region are denoted as *ℓ*_1_ and *ℓ*^2^, which are described by *α*_A_=*α*_l_*α*_p53_ and *α*_A_=*α*_2_*α*_p53_, where a and a are all positive constants satisfying *α*_1_, the timing variability below *ℓ*_1_ or beyond *ℓ*_2_, is less than that in the strip region, and the timing variability below *ℓ*_1_ is less than that beyond *ℓ*_2_, (referring to the dashed line with arrow). These indicate that in the outside of the strip region, if the timescale of protein *A* is dominant, the timing variability becomes smaller, and conversely, if the timescale of protein *p*53 is dominant, the timing variability also becomes smaller. Since timescale factors *α*_p53_ and *α*_A_ determine the noise in proteins *p*53 and *A* (called intrinsic and extrinsic noise) respectively, we can thus conclude that the intrinsic and extrinsic noise all can significantly contribute to the timing variability, but this contribution depends on which noise is dominant. This is an interesting phenomenon similar to the resonance that takes place as the external frequency is approximately equal to the external frequency (36).

**FIGURE 4.**
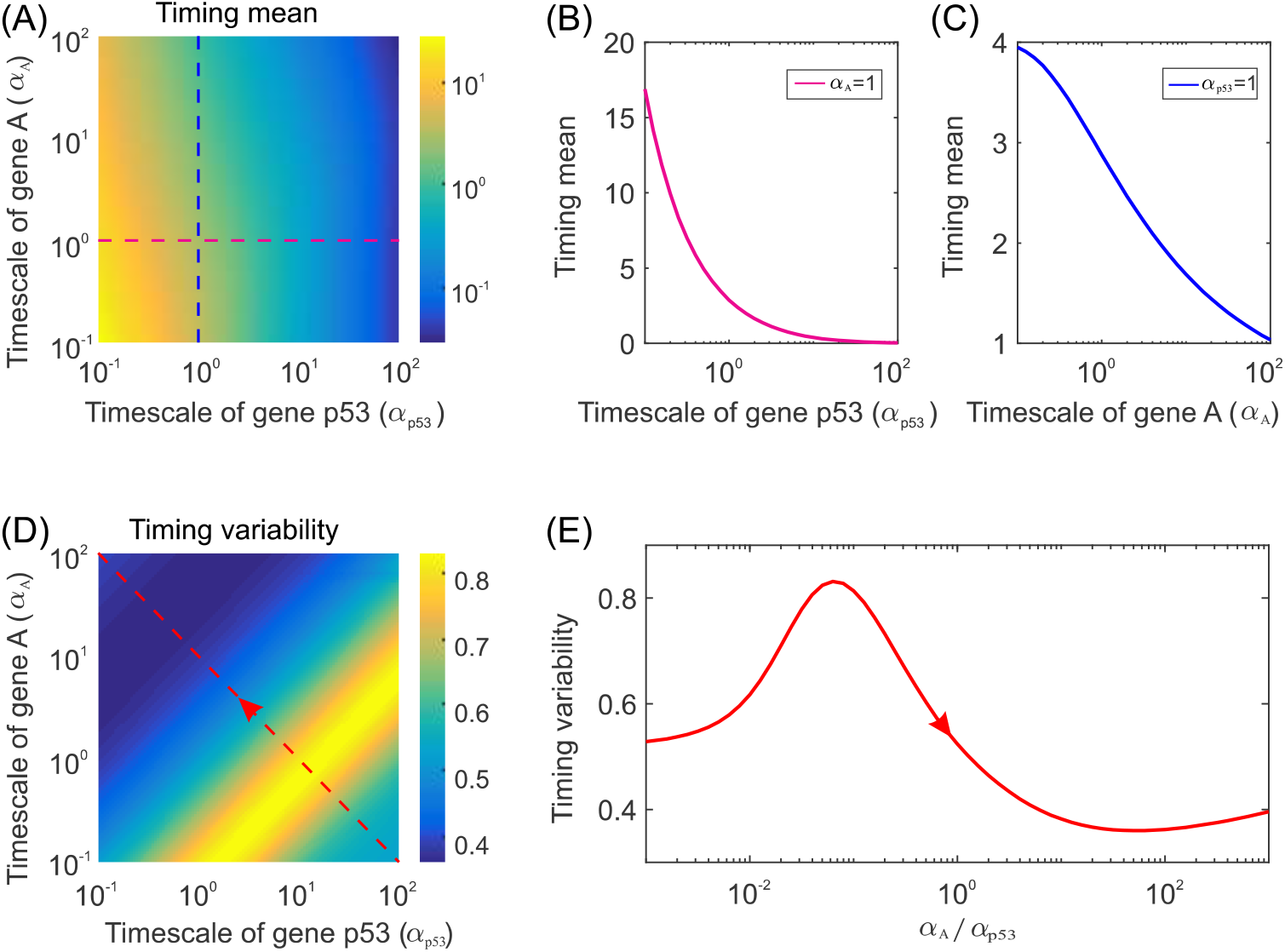
**Effects of timescales on the timing of events**, where *α*_p53_ and *α*_A_ are respectively the common factors that the transcription and degradation rates of proteins *p*53 and *A* are simultaneously enlarged. (A) Heatmap showing timing mean as a function of *α*_p53_ and ^α^_A_. (B) Timing mean as a function of the timescale of protein *p*53, where *α*_A_= 1. (C) Timing mean as a function of the timescale of protein *A*, where *α*_p53_= 1. (D) Heatmap showing timing variability as a function of the timescales of protein *p*53 and *A*. (E) Timing variability as a function of the rate of the timescale of protein *A* over that of protein *p*53, where arrow is in agreement with that in (D). In (A)-(E), the parameter values are *g*_p53_= 15, *d*_p53_= 1, *b*= 1, *g*_A_= 15, *d*_A_ = 1 and *p*53_max_ = 50.

From Figs. 4*B* and 4*C*, we observe that the mean FPT is a monotonically decreasing function of the timescale for proteins *A* and *p*53 respectively, but the former is a convex-downward curve for a particular timescale of protein *p*53, i.e., for *α*_A_= 1, whereas the latter is fundamentally a line for a particular timescale of protein *A*, i.e., for *α*_p53_= 1. Note that *α*_p53_= 1 and *α*_A_= 1 correspond to black and red dashed lines in Fig. 4*A*. The results shown in Figs. 4*B* and 4*C* are practically special results shown in Fig. 4*A*. Figures 4*B* and 4*C* imply that internal and external (or threshold) timescales can all shorten the mean arriving time (i.e., the mean threshold crossing time). This further implies that timescales can speed up response.

Figure 4*E*, which corresponds to the red dashed with arrow inn Fig. 4D, shows how the rate between the timescales of proteins *p*53 and *A*, *γ*=*α*_A_/*α*_p53_, impacts the variability in the event timing. Interestingly, we observe that there is an optimal rate between external and internal timescales, *γ*_critical_, such as the timing variability is maximal, implying that the precision in the event timing is worst for this optimal timescale. Furthermore, the rate *γ* satisfying *γ*<*γ*_mitical_ increases the timing variability whereas the rate *γ* satisfying *γ*>γ_critical_ fundamentally decreases the timing variability. The former implies that when the rate of the external timescale over the internal timescale, *γ*, is less than the critical rate, *γ*_critical_, this rate weakens the precision in the event timing, and conversely, it fundamentally enhances this precision.

In a word, the timescales of proteins *p*53 and *A* are two unneglectable factors in the timing of events since they can significantly affect the timing precision and the arriving time (or the mean FPT). And there is a strip region in the plane of external and internal timescales such that the timing variability is largest (implying that the timing precision is worst).

### Effects of p53 transcription and degradation rates on fractional killing of cancer cells

In our model, apart from parameters associated with promoter kinetics, there are the protein transcription and degradation rates involved. The curves shown in Fig. 4 correspond to a special value of the transcription or degradation rate of protein. However, these two rates can be regulated by external signals, leading to changes in biologically reasonable intervals. This raises a question: how the two parameters impact the arriving time and the variability in the event timing. Here we numerically analyze this impact, with results shown in Fig. 5.

**Fig. 5.**
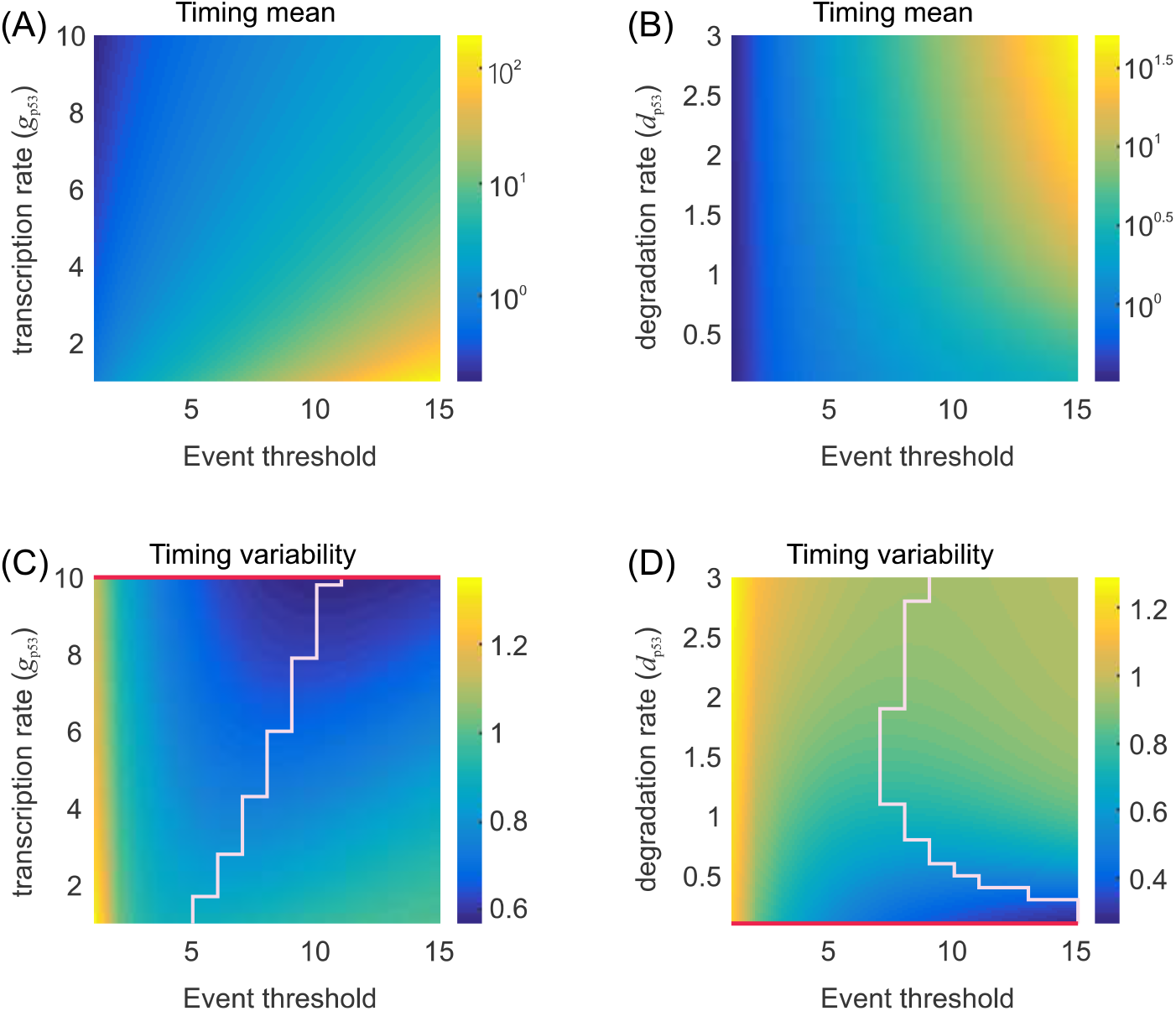
Influence of transcription or degradation rate on the timing of events. **(A)** Heatmap showing timing mean as a function of both event threshold and transcription rate (*g*_p53_). (B) Heatmap showing timing mean as a function of event threshold and degradation rate (*d*_p53_). (C) Heatmap showing timing variability as a function of event threshold and transcription rate (*g*_p53_), where the stair-like line consists of the points corresponding to the star in Fig. 3(D) where in a special value of *g*_p53_ is used. (D) Heatmap showing timing variability as a function of event threshold and degradation rate (*d*_p53_), where the meaning of the stair-like line is similar to that in (C). Note that the event of threshold crossing has taken place when *d*_p53_ is sufficiently small, which corresponds to the red line. In (A) -(D), the parameter values are set as *b* = 1, *g*_A_=10, *d*_A_ =, *p*53_max_ = 30 and *A*_thresold_ = 1-15. In (A) and (C), we consider the range of g_p53_ = 1 ~ 10 and *d*_p53_ = 1, whereas in (B) and (D), we set *g*_p53_= 5 and *d*_p53_ = 0.1 ~ 3.

From Fig. 5*A*, we observe that for a fixed event threshold, the mean FPT monotonically decreases with increasing the transcription rate (*g*_p53_), implying that the transcription rate of protein *p*53 shortens the arriving time or accelerates the threshold crossing. On the other hand, for a fixed transcription rate (*g*_p53_), the mean FPT is a monotonically increasing function of event threshold, implying that the event threshold slows down the threshold crossing. By comparing Fig. 5B with Fig. 5*A*, we can see that the change tendency in the case of degradation rate is fundamentally opposite to that in the case of transcription rate.

From Fig. 5C, we observe that for a fixed event threshold, the variability in the timing is monotonically decreasing with the increase of transcription rate (*g*_p53_). We also observe from this panel that for a fixed yet small transcription rate (*g*_p53_), the variability in the timing first decreases and then increases with the increase of event threshold. Fig. 5C also shows a stair-like line, which is composed of the points corresponding to the least timing variability. From Fig. 5D, we also observe that for a fixed degradation rate (*d*_p53_), there exists a minimal timing variability. But for a fixed event threshold, the mean FPT is a monotonically increasing function of the degradation rate (*d*_p53_).

In a short, both smaller transcription rates and larger event thresholds or both larger degradation rates and larger event thresholds lead to larger mean FPTs, implying that fewer cancer cells are killed. However, the results in the case of timing variability is almost converse to those in the case of mean FPT.

### Effect of burst size in p53 on fractional killing of cancer cells

Here we investigate the influence of burst size on mean FPT and timing variability, with results shown in Fig. 6. From Fig. 6*A*, we observe that the mean FPT is a monotonically decreasing function of mean burst size. Moreover, the curve corresponding to fluctuating threshold is always below the curve corresponding to fixed threshold, implying that dynamically fluctuating thresholds accelerate threshold crossing. Fig. 6*B* shows the dependence of variability in timing on the mean burst size (*b*) for two different cases: fixed and fluctuating thresholds.

**FIGURE 6.**
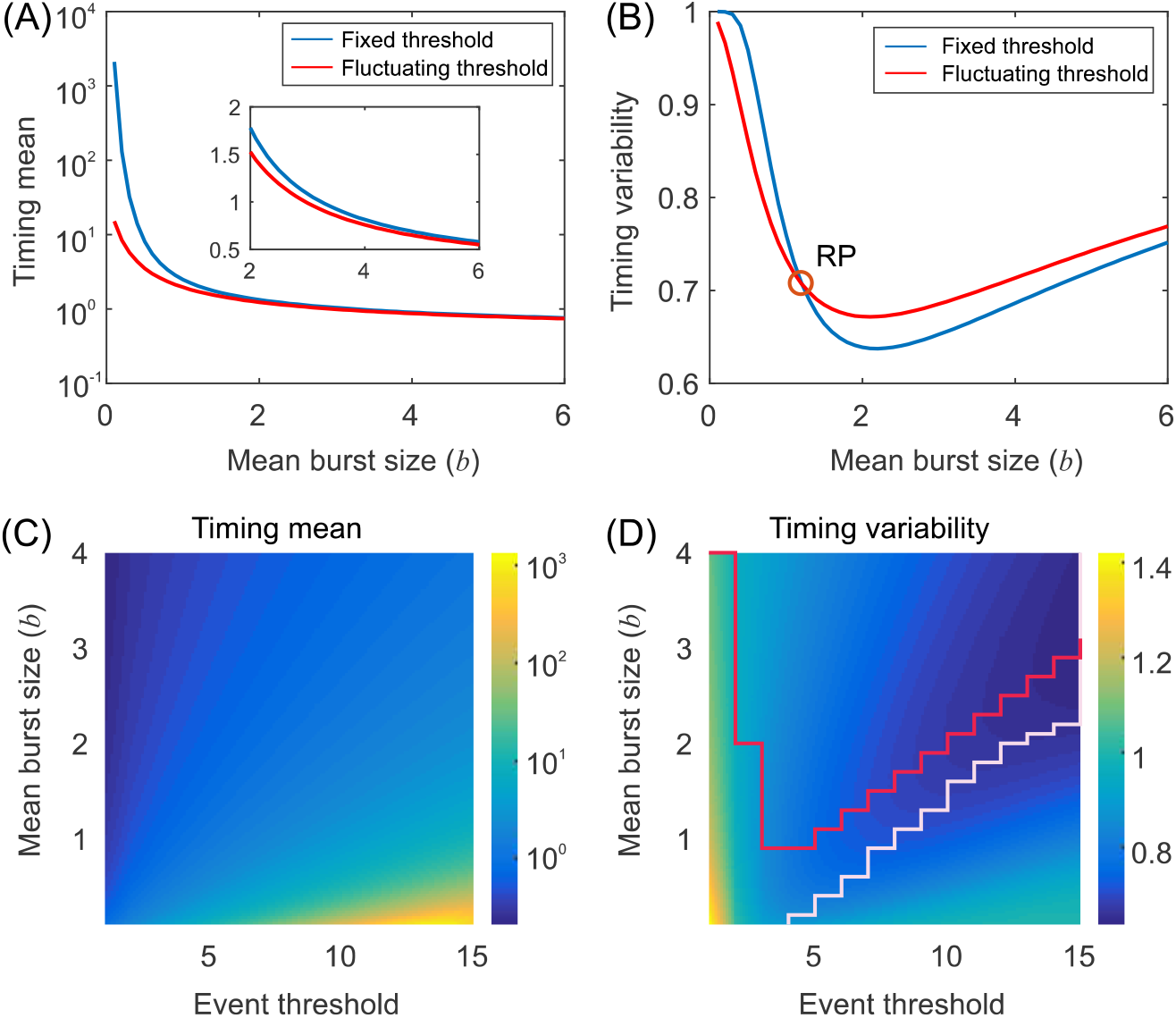
Effects of mean burst sizes on event timing. (A) Timing mean as a function of the mean burst size, where the inset shows a partial enlarged diagram. (B) Timing variability as a function of the mean burst size two different cases: fixed threshold (blue curve) and fluctuating threshold (red curve), where RP represents a critical point for reversing. (C) Heatmap showing timing mean as a function of event threshold and mean burst size. (D) Heatmap showing timing variability as a function of event threshold and mean burst size, where the red curve corresponds to the case that an event of threshold crossing has happened either due to a sufficiently large transcription rate (*g*_p53_) or due to a sufficiently small degradation rate (*d*_p53_), and the stair-like line consists of the points corresponding to the star in Fig 3(C) wherein a special value of g_p53_ is used. In (A)-(D), the parameter values are set as *g*_p53_= 5, *d*_p53_= 1, *g_A_* = 10 and *d_A_*, = 1. The cutoff constant is set as *p*53_mav_ = 30. In (A) and (B), the range of mean burst size is set as *h*= 0.1 – 6 and the threshold is set as *A*_thresold_=10, whereas in (C) and (D), the range of mean burst size is set as *b* = 0.1 ~ 4 and the range of threshold is set as *A*_thresold_ = 1 ~ 15.

From Fig. 6*B*, we observe that there is optimal *b* such that the variability in the timing is least in both cases of threshold. This implies the mean burst size can make the timing precision reach optimality in both cases of fixed and fluctuating thresholds. In addition, we observe that there is a critical *b* (denoted by RP) such that the timing variability in the case of fluctuating threshold is smaller than that in the case of fixed threshold, as *b* is below RP but the former is larger than the latter as the event threshold is beyond RP. In other words, for a small *b*, threshold fluctuations can reduce the timing variability or raise the precision in the timing.

From Fig. 6C, we observe that larger mean FPTs appear in the region of the right-down corner, which corresponds to both smaller mean burst size and larger event thresholds. Specifically, for a fixed mean threshold, a larger mean burst size leads to the reduction of the mean FPT, implying that translation burst can accelerate response by shortening the arriving time. On the other hand, for a fixed mean burst size, a larger mean threshold leads to the increase of the mean FPT, implying that fluctuating threshold can slow down response by prolonging the arriving time. From Fig. 6*D* (red curve), we observe that larger timing variability appears approximately in the region of the left-down corner, which corresponds to both smaller mean burst size and smaller event thresholds. Specifically, for a fixed small mean threshold, a smaller mean burst size leads to the increase of timing variability, implying that translation burst can slow down response by increasing the timing variability. However, there exists an optimal mean burst size such that the timing precision is best for almost large mean threshold. On the other hand, for a fixed large mean burst size (referring white curve in Fig. 6*D*), a larger mean threshold leads to the decrease of the timing variability, implying that fluctuating threshold can enhance the timing precision by reducing the timing variability. However, there exists an optimal mean threshold such that the timing precision is best for almost large mean.

In short, translation burst (internal noise) accelerates threshold crossing (implying that more cancer cells are killed), there is a critical mean burst size such that translation burst enhances the timing precision as the mean burst size is below this critical value but reduces the timing precision as the mean burst size is beyond this critical value, and there is an optimal mean burst size such that the timing precision is best.

## Conclusion and Discussion

While fractional killing is a major impediment to the treatment of cancer, viruses, and microbial infections, non-genetic variability plays a pivotal role in fractional killing. Sources of this variability may be complex: apart from molecular noise inherent to gene expression, there is dynamic variability in apoptotic threshold (2,3). In this paper, we have systematically investigated a stochastic gene expression system underlying the process of fractional killing, where the cell is killed only when the p53 expression level crosses a dynamically fluctuating threshold for the first time. The main contributions and insights can be summarized as follows: (1) fluctuations in apoptotic threshold accelerate response, and a faster fluctuation leads to a smaller mean FPT or to killing of more cancer cells; (2) there is an optimal mean threshold such that the timing variability is least; (3) there is an optimal mean burst size such that the timing repression is best or the timing variability is smallest; (4) for a high enough threshold, fluctuations in threshold can raise the timing precision; and (5) the timescales between transcription and degradation rates can adjust the precision in the timing, independent of the ratio of the transcription rate over the degradation rate. These results indicate that in contrast to fixed apoptotic thresholds, fluctuating apoptotic thresholds can significantly influence the timing of events or killing of cancer cells.

Although we used a simple stochastic model to investigate fractional killing processes involving timing events, our theoretical framework, i.e., a one-dimensional FPT problem with a dynamically fluctuating boundary is transformed into a two-dimensional FPT problem with a fixed boundary can easily extended to other complex or general cases. In fact, timing events can be attributed to a canonical mechanism of threshold crossing (that can occur in many cellular processes ranging from responses of cells to their environmental cues to cell cycles and circadian clocks), by which a molecular event triggers a cellular behavior is accumulation to a threshold (17,18,37–40). In this mechanism, molecules are steadily produced by the cell, and once the molecule number crosses a particular threshold, the behavior is initiated. Most of these threshold-crossing processes are based on gene expression, e.g., an activated gene may be required to reach in a precise time a threshold level of gene expression that triggers a specific downstream pathway. However, a gene may reach a critical threshold of expression with substantial cell-to-cell variability even among isogenic cells exposed to the same constant stimulus. This variability is a necessary consequence of the inherently stochastic nature of gene expression (41–47). Apart from this internal stochastic origin of timing, dynamically fluctuating thresholds can also result in variability in the timing required to reach a critical threshold level. It is possible that the intrinsic ‘molecular noise’ in intracellular processes is responsible for such cell-to-cell variability in timing. This is experimentally difficult to verify, but may beg theoretical analysis as done in this paper.

How robust are our results to noise sources and key modeling assumptions? For example, our model only considers the intrinsic noise in gene product levels but ignores the extrinsic noise in gene expression machinery (48,49). To incorporate such extrinsic noise, one may alter the transcription rate to *k_i_Z*(*k_i_* is an external parameter), where *Z* may be drawn from an a priori probability distribution at the start of gene expression (*t*= 0) and remains fixed till the threshold is reached. Our model also ignored feedback regulation, which however exists widely in biological regulatory systems. Recent work has investigated the impact of feedback regulation on the timing of events in the case of fixed threshold (17,18,40). Interestingly, it was found that there is an optimal feedback strategy to regulate the synthesis of a protein to ensure that an event will occur at a precise time, while minimizing deviations or noise about the mean. In spite of this, how feedback regulation controls or impacts the timing of events in the case of dynamically fluctuating threshold is unclear. Using our analysis framework, one can also study the effect of feedback regulation on the timing of events in the case of dynamically fluctuating threshold. In our case, if changes in burst size, transcription rate or degradation rate are taken as the consequence of feedback regulation, the effect of feedback regulation on the timing of events will become clear in the case of fluctuating threshold. In addition, complex regulatory network composed of apoptosis related proteins also plays an important role in fractional killing. It is promising for future work to study how cross talks between the apoptosis pathway and survival pathways affect fractional killing (50).

Next, we simply discuss potential biological implications of our results in the context of fractional bacterial killing and p53 dynamics.

### Connecting theoretical insights to fractional killing

Exposure of an isogenic bacterial population to a cidal antibiotic typically fails to eliminate a small fraction of refractory cells. In order to interpret this phenomenon, Roux J, et al. (5) investigated the basis of fractional cell killing by TRAIL and antibody agonists of DR4 and DR5 receptors. They demonstrated the existence of a threshold in initiator caspase activity (referred to as C8) that must be exceeded for cells to die. Interestingly, they found that in cells that go on to die, C8 activity rises rapidly and monotonically until the threshold is reached and mitochondrial outer membrane permeabilization ensues, whereas in cells that survive, C8 activity rises more slowly for 1-4 h, never achieving the level required for death, and then falls back to pre-treatment levels over the next 4-8 h due to proteasome-mediated protein degradation. This finding, which can be reproduced by analysis of our model through the proposed method, implies that Mycobacterium smegmatis can dynamically persist in the presence of a drug, and the stable number of cells characterizing this persistence was actually a dynamic state of balanced division and death.

### Connecting theoretical insights to drug therapy

Many chemotherapeutic drugs kill only a fraction of cancer cells, limiting their effectiveness. Paek AL, et al. (3) used live cell imaging to check the role of p53 dynamics in fractional killing of colon cancer cells in response to chemotherapy. They found that both surviving and dying cells reach similar levels of p53, indicating that cell death is not determined by a fixed p53 threshold. Instead, a cell’s death probability relies on the time and levels of p53. Cells must reach a critical threshold level of p53 to execute apoptosis, and this threshold increases over time. The increase in p53 apoptotic threshold is due to drug-dependent induction of anti-apoptotic genes, predominantly in the inhibitors of apoptosis family. While that study underlined the importance of measuring the dynamics of key players in response to chemotherapy to determine mechanisms of resistance and optimize the timing of combination therapy, our study here provided quantitative results for this importance.

Finally, from a theoretical point of view, our work provides a mathematical and computational framework for studying how fluctuations in threshold influence the statistics of FPT. And our methods can be extended to the analysis of fluctuations of derivative thresholding (5), integral thresholding (51), oscillation (52). Exploring these constraints in more detail will be an important avenue for future research. In addition, analytical results and insights obtained here have broader implications for timing phenomenon in chemical kinetics, epidemic spreading, ecological modeling, and statistical physics. In addition, our methods may allow us to better understand the complex patterns of sequentially ordered biochemical events that are often observed in development and cell-fate decision presumably require an effective control of event timing (53–57).

## Supporting information

S1 Supporting Information. Mathematical derivations and supplementary information.

## Supporting Information

S1 Supporting Information. Mathematical derivations and supplementary information. Text

S1. The derivation of all equations in the main text is provided, and more numerical results are demonstrated.

## Acknowledgements

This work was supported by grants 91530320, 11775314, 11475273, and 11631005 from Natural Science Foundation of P. R. China; 2014CB964703 from Science and Technology Department, P. R. China; 201707010117 from the Science and Technology Program of Guangzhou, P. R. China.

## Author Contributions

Conceived and designed the experiments: JJ, TS. Performed the experiments: BH, JJ. Analyzed the data: BH, JJ, TS. Contributed reagents/materials/analysis tools: JJ, TS. Wrote the paper: JJ, TS.

