## Supplementary material for "Dynamic variability in apoptotic threshold as a strategy for combating fractional killing": S1 Supporting Information. Mathematical derivations and supplementary information.

### Dynamic variability in apoptotic thresholds as a strategy for combating fractional killing of cancer cells

1. Guangdong Province Key Laboratory of Computational Science, Sun Yat-Sen University, Guangzhou, People's Republic of China.
2. School of Mathematics, Sun Yat-Sen University, Guangzhou, People's Republic of China

In this supplement, we provide details for mathematical derivations in the main text and present figures that illustrates some of the results in greater depth.

### 1 Mathematical modeling for FPT problem

#### 1.1 A FPT problem with fluctuating threshold

Let $\{p53: p53 = p53(t), t \geq 0 | p53(0) = p53_0\}$ be a temporally homogeneous diffusion process, and $\{A: A = A(t), t \geq 0 | A(0) = A_0\}$ be a fluctuating barrier (called the barrier system). Without loss of generality, we set $p53_0 < A_0$ or $p53(0) < A(0)$. This setting is natural since $A$ represents a boundary or threshold that $p53$ will cross. Note that the union of $p53$ and $A$, $\{(p53, A): p53 = p53(t), A = A(t), t \geq 0 | p53(0) = p53_0, A(0) = A_0\}$ constitutes a new system or a new process. Define $T$ as the time that $p53$ hits the fluctuating barrier $A$ for the first time, that is (1, 2, 3),

$$T = \min\{t: p53(t) \geq A(t) | p53(0) = p53_0 < A(0) = A_0\} \quad \text{(S1)}$$

which is called the first passage time (FPT). Apparently, $T$ is a random variable since both $p53(t)$ and $A(t)$ are random, referring to Fig. S1*A*, and depends on $p53_0$ and $A_0$.

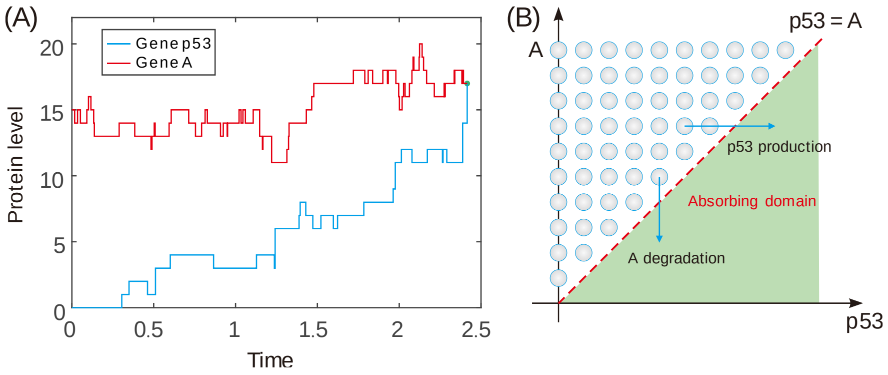

**Fig. S1 Event timing is modeled as a problem of first passage time (FPT).** (A) shown is an example for FPT, where $A(t)$ represents the boundary and the solid circle represents the point that stochastic trajectory $p53(t)$ hits stochastic trajectory $A(t)$ for the first time; (B) an absorbing domain of PFT, defined by $\mathcal{D} = \{(p53, A) | p53 \geq A\}$. Note that the timing event is triggered once $p53$ reaches $A$.

Then, we define an absorbing domain $\mathcal{D}$, which consists of those points $(p53, A)$ that satisfy $p53 \geq A$ in the $(p53, A)$ plane, that is, $\mathcal{D} = \{(p53, A) | p53 \geq A\}$. In addition, we define a survival probability, $\mathbb{S}$, that the trajectory starting from $\{(p53(t), A(t))\}$ at time $t=0$ has not yet been absorbed into domain $\mathcal{D}$ at time $t$. Let $P_{X',Y'}(t)$ represent the probability that the system $\{(p53, A)$: $p53 = p53(t), A = A(t), t \geq 0 | p53(0) = p53_0 < A(0) = A_0\}$ is at state $(X', Y')$ at time $t$, that is,

$$P_{X',Y'}(t) = \text{Prob}\{p53(t) = X', A(t) = Y' | p53(0) = p53_0 < A(0) = A_0\} \tag{S2}$$

For convenience, $P_{X',Y'}(t)$ is sometimes denoted by $P_{S'}(t)$, i.e., $P_{S'}(t) = P_{X',Y'}(t)$, where $S' = (X', Y')$ represents state. Note that the survival probability can be expressed as

$$\mathbb{S} = \sum_{S \neq D} P_S(t). \tag{S3}$$

Denote by $f_T(t)$ the probability density function of the FPT, that is,

$$f_T(t) = \text{Prob}\{T \leq t\}. \tag{S4}$$

In the following, we are mainly interested in statistical properties of random variable $T$. For this, we first establish the relation between $f_T(t)$ and $\mathbb{S}$, and then give the expression of $f_T(t)$.

#### 1.2 Probability density function of FPT

Assume that all states $\{S(t): t \geq 0\}$ with $S(t) = (p53(t), A(t))$ constitute a Markov process. Then, we have a forward master equation (FME) of the form

$$\frac{\partial \mathbf{P}(t)}{\partial t} = \mathbf{MP}(t) \tag{S5}$$

where $\mathbf{P}(t)$ is a column vector consisting of all $P_S(t)$, and $\mathbf{M}$ is a certain linear operator, depending a process of interest. Note that every component of $\mathbf{P}(t)$ is the probability that the system $\{S(t): t \geq 0\}$ arrives at the absorbing domain $\mathcal{D}$ at time $t$, and that $\mathbf{M}$ is actually a state transition matrix, whose components can be expressed as

$$\mathcal{M}_{S,S'} = \alpha(S, S') - \delta_{S,S'} \sum_{S''} \alpha(S'', S) \tag{S6}$$

where $\alpha(S, S')$ denotes the transition rate from state $S'$ to state $S$, and $\delta_{S,S'}$ is the Kronecker delta function.

On the other hand, there is an absorbing state $S_f$ with $S_f \in \{(p53_f, A_f)\}$, such that the transition rate from absorbing state $S_f$ to state $S$ is always zero, that is, $\alpha(S,S_f)=0$ for all states $S$. According to the definition of $\mathcal{M}_{S,S'}$, we can know the conservative condition: $\sum_S \mathcal{M}_{S,S'}=0$. Let $P(S_f,t|S,t_0)$ be the probability that the state $S(t)$ reaches the absorbing state $S_f$ at time $t$, given the initial state $S=S(t_0)$ at time $t_0$ with $S(t_0)=(p53_0,A_0)$ (usually, $t_0=0$ is set). Then, we can determine the probability density function, $f(t;S_f|S)$, of the FPT for state $S$ to reach absorbing state $S_f$.

Denote by $\mathbb{S}(t,S_f|S,t_0)$ the survival probability that the trajectory $S(t)$ starting from $S$ at time $t_0$ has not yet been absorbed to state $S_f$ at time $t$, that is,

$$\mathbb{S}(t,S_f|S,t_0)=\sum_{S'\neq S_f} P(S',t|S,t_0) \tag{S7}$$

By definition, we can have

$$\text{Prob}\{T\le t\}=1-\mathcal{S}(t,S_f|S,t_0). \tag{S8}$$

In fact, the probability that state $S$ reaches state $S_f$ in an infinitesimal time interval $(t,t+dt)$ is $f(t;S_f|S)dt$, so the probability that state $S$ reaches state $S_f$ at time $t$ is given by $\int_0^t f(t';S_f|S)dt'$. Thus, the survival probability $\mathbb{S}(t,S_f|S,t_0)$ is related to the FPT distribution through the following manner

$$\mathbb{S}(t,S_f|S,t_0)=1-\int_0^t f(t';S_f|S)dt' \tag{S9}$$

In turn, we can determine the probability density function of the FPT. In fact, it follows directly from Eq. S9 that

$$f(t;S_f|S)=-\frac{\partial\mathbb{S}(t,S_f|S,t_0)}{\partial t} \tag{S10}$$

Using Eq. S5 and Eq. S7, we further have (1, 2)

$$\begin{aligned}\partial_t\mathbb{S}(t,S_f|S,t_0)&=\partial_t\sum_{S'\neq S_f}P(S',t|S,t_0)=\sum_{S'\neq S_f}\partial_t P(S',t|S,t_0)\\&=\sum_{S'\neq S_f}\sum_{S''\neq S_f}\mathcal{M}_{S',S''}P(S'',t|S,t_0)=\sum_{S''\neq S_f}P(S'',t|S,t_0)\sum_{S'\neq S_f}\mathcal{M}_{S',S''}\\&=\sum_{S''\neq S_f}P(S'',t|S,t_0)\left(\sum_{S'}\mathcal{M}_{S',S''}-\mathcal{M}_{S_f,S''}\right)=-\sum_{S''\neq S_f}\mathcal{M}_{S_f,S''}P(S'',t|S,t_0)\end{aligned} \tag{S11}$$

where we have used the fact of $\sum_{S'}\mathcal{M}_{S',S''}=0$ due to the probability conservation. Thus, the probability density function of the FPT is given by

$$f_T(t)\equiv f(t;S_f\,|\,S)dt=\sum_{S''\neq S_f}\mathcal{M}_{S_f,S''}P\left(S'',t\,|\,S,t_0\right) \tag{S12}$$

which can simply be expressed as the following form of vector

$$f_T(t)=\mathbf{W}^{\mathrm{T}}\mathbf{P}(t) \tag{S13}$$

Note that Eq. S13 has the solution of the form

$$f_T(t)=\mathbf{W}^{\mathrm{T}}\exp(\mathbf{M}t)\mathbf{P}(0) \tag{S14}$$

where $\mathbf{W}$ is the column vector of the transition rates from all accessible states $S''$ to state $S_f$.

##### 1.3 Statistical quantities of FPT

Once the probability density function of the FPT $f_T(t)$ is given, we can calculate raw moments of random variable $T$, according to

$$\left\langle T^k\right\rangle=\int_0^{+\infty}t^k f_T(t)dt\,,\quad k=1,2,\cdots \tag{S15}$$

Substituting the expression of $f_T(t)$ into the above Eq. S15, and using the integration by parts for many times, $\left\langle T^k\right\rangle$ can be transformed the following form

$$\begin{aligned}\left\langle T^k\right\rangle&=\mathbf{W}^{\mathrm{T}}\left[\int_0^{\infty}t^k\exp(\mathbf{M}t)dt\right]\mathbf{P}(0)\\&=\mathbf{W}^{\mathrm{T}}\left[t^k\mathbf{M}^{-1}\exp(\mathbf{M}t)\Big|_0^{\infty}-k\mathbf{M}^{-1}\int_0^{\infty}t^{k-1}\exp(\mathbf{M}t)dt\right]\mathbf{P}(0)\\&=(-1)^{k+1}k!\,\mathbf{W}^{\mathrm{T}}\left(\mathbf{M}^{-1}\right)^{k+1}\mathbf{P}(0)\end{aligned} \tag{S16}$$

where the characteristic values of matrix $\mathbf{M}$ are assumed to have negative real parts, and $t^k\mathbf{M}^{-1}\exp(\mathbf{M}t)$ $(k=0,1,2,\ldots)$ in the above expression goes to zero. Moreover, $\mathbf{W}^{\mathrm{T}}\mathbf{M}^{-1}=-\mathbf{e}^{\mathrm{T}}$ can be obtained due to $\langle T\rangle\equiv 1$, where $\mathbf{e}^{\mathrm{T}}=[1,1,\cdots,1]$ is a constant vector. Thus we easily show

$$\left\langle T^k\right\rangle=\int_0^{+\infty}t^k f_T(t)dt=k!(-1)^k\mathbf{e}^{\mathrm{T}}\left(\mathbf{M}^{-1}\right)^k\mathbf{P}(0),\quad k=1,2,\cdots \tag{S17}$$

Furthermore, it follows from Eq. S17 that the mean first passage time (MFPT) of the probability density function $f_T(t)$ and the intensity of the noise with $T$ of FPT, (defined as the ratio of variance over the square of mean), denoted by $\mathrm{CV}_T$, are given by

$$\mathrm{MFPT}=\langle T\rangle=-\mathbf{e}^{\mathrm{T}}\mathbf{M}^{-1}\mathbf{P}(0), \tag{S18}$$

$$\mathrm{CV}_T = \frac{\langle T^2\rangle - \langle T\rangle^2}{\langle T\rangle^2} = \frac{2\mathbf{e}^{\mathrm{T}}\left(\mathbf{M}^{-1}\right)^2\mathbf{P}(0)}{\left[\mathbf{e}^{\mathrm{T}}\mathbf{M}^{-1}\mathbf{P}(0)\right]^2} - 1. \tag{S19}$$

In the following, "Timing Mean" and "Timing variability" are measured by MFPT and $\mathrm{CV}_T$, respectively.

In addition, we can calculate the high order features, such as skewness and kurtosis, according to calculation formula

$$\mathrm{Skew}_T = \frac{\langle T^3\rangle - 3\langle T\rangle\langle T^2\rangle + 2\langle T\rangle^3}{\left(\langle T^2\rangle - \langle T\rangle^2\right)^{3/2}} \tag{S20}$$

$$\mathrm{Kurt}_T = \frac{\langle T^4\rangle - 4\langle T\rangle\langle T^3\rangle + 6\langle T\rangle^2\langle T^2\rangle - 3\langle T\rangle^4}{\left(\langle T^2\rangle - \langle T\rangle^2\right)^2} \tag{S21}$$

We can give the explicit expressions of skewness and kurtosis, based on Eq. S17.

### 2 Gene expression model of FPT with fluctuating threshold

From now on, we consider a common model of stochastic gene expression with a stochastically fluctuating threshold, referring to Fig. S2. A threshold event is triggered once the expression level of a gene, denoted by $p53$, crosses the expression level of another gene, denoted by $A$ (as a critical threshold) for the first time. We are interested in the stochastic but not deterministic threshold crossing.

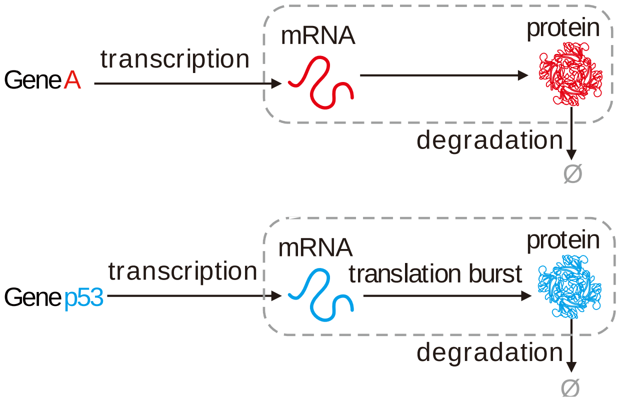

**Fig. S2 Schematic for a minimal model of FPT**, where a threshold event is triggered once the expression level of a gene crosses the expression level of another gene (as a critical threshold) for the first time.

Assume that the $p53$ molecules are produced in a burst manner whereas the $A$ molecules are generated in a constitutive manner. We use the produced counts of protein $A$ to construct a stochastically fluctuating threshold that the molecule number of protein $p53$ reaches. Let

$A(t) \in \{0,1,2,\cdots\}$ denote the level of protein $A$ at time $t$, and assume that $A(t)$ follows a Poisson distribution with two characteristic parameters $g_{\mathrm{A}}^{(n)}$, (where superscript $(n)$ means that feedback regulation is considered. This superscript may be omitted in the absence of feedback regulation) and $d_{\mathrm{A}}$ representing respectively the transcription and degradation rates of protein $A$ when $A(t)=n$. Let $p53(t) \in \{0,1,2,\cdots\}$ denote the level of protein $p53$ at time $t$, and assume that protein $p53$ is generated with a Poisson rate $g_{\mathrm{p53}}^{(m)}$ (where superscript $(m)$ means that feedback regulation is considered, and it may be neglected in the absence of feedback regulation), and degrades at a constant degradation rate $d_{\mathrm{p53}}$. The translation burst approximation is based on the assumption of short-lived mRNAs, that is, each mRNA degrades instantaneously after producing a burst of $B$ protein molecules, where $B$ follows a geometric distribution assumed as usual

$$P_B(B=k) = \frac{b^k}{(1+b)^{k+1}}, k = 0,1,2,\cdots, b > 0 \quad \text{(S22)}$$

Then, we can show

$$P_B(B \geq k) = \left(\frac{b}{1+b}\right)^k, k = 0,1,2,\cdots \quad \text{(S23)}$$

where $b$ represents the mean protein burst size. In what follows, we denote $P_{B=k} \equiv P_B(B=k)$ and $P_{B\geq k} \equiv P_B(B \geq k)$ for convenience.

The time evolution of $(p53(t), A(t))$ starting from $p53(0)=m, A(0)=n$ with $m<n$ at time $t=0$ can be descried through the following probabilities of timing events in the next infinitesimal time $(t, t+dt)$

$$\begin{aligned}
&P\left(p53(t+dt)=m+B, A(t+dt)=n \middle| p53(t)=m, A(t)=n\right) = g_{\mathrm{p53}}^{(m)}dt;\\
&P\left(p53(t+dt)=m-1, A(t+dt)=n \middle| p53(t)=m, A(t)=n\right) = md_{\mathrm{p53}}dt;\\
&P\left(p53(t+dt)=m, A(t+dt)=n+1 \middle| p53(t)=m, A(t)=n\right) = g_{\mathrm{A}}^{(n)}dt;\\
&P\left(p53(t+dt)=m, A(t+dt)=n-1 \middle| p53(t)=m, A(t)=n\right) = nd_{\mathrm{A}}dt;
\end{aligned} \quad \text{(S24)}$$

An event occurs if the cumulative number of proteins $p53$ reaches the number of protein $A$ molecules. The relation between the proteins $p53$ and $A$ can be considered as a trajectory in the domain $\{(p53, A) | p53 = p53(t) < A(t) = A\}$. The corresponding forward master equation (FME) describing the time evolution of protein pair $p53$ and $A$ can be described as [4]

$$\frac{dP_{m,n}(t)}{dt} = \sum_{i=0}^{m-1} g_{\text{p53}}^{(i)} P_B(B=m-i) P_{i,n}(t) + g_{\text{A}} P_{m,n-1}(t) + (m+1) d_{\text{p53}} P_{m+1,n}(t) + (n+1) d_{\text{A}} P_{m,n+1}(t) - \left(g_{\text{p53}}^{(m)} P_B(B \geq 1) + g_{\text{A}} + m d_{\text{p53}} + n d_{\text{A}}\right) P_{m,n}(t) \tag{S25}$$

where $m < n$. And the probability density function of the FPT (or the FPT distribution ) can be formally expressed as

$$f_T(t) = \sum_{m \geq 0} d_{\text{A}} (m+1) P_{m,m+1}(t) + \sum_{n > m \geq 0} g_{\text{p53}}^{(m)} P_B(B \geq n-m) P_{m,n}(t) \tag{S26}$$

In what follows, we will consider only the case that there is no feedback and the transcription rates are constants, i.e., $g_{\text{p53}}^{(m)} \equiv g_{\text{p53}}$ and $g_{\text{A}}^{(n)} \equiv g_{\text{A}}$. To determine the first several moments of the FPT to a fluctuating threshold, we need the specific expression of the PFT distribution $f_T(t)$.

### 3 FPT distribution and its statistics in four different fluctuating thresholds

For clarity, we will distinguish four cases of the absorbing domain to solve the FPT problem formulated above. These cases are schematized in Fig. S3. Based on the finite state projection approach, we construct a new process for pair $p53$ and $A$ on the finite state-space

$$\Omega = \left\{ \left( p53(t), A(t) \right) \middle| \, p53(t) = 0,1,2,\cdots,C_{\text{p53}}, A(t) = 0,1,2,\cdots,C_{\text{A}} \right\}, \tag{S27}$$

where the states represent the numbers of proteins $p53$ and $A$. Here we introduce two numerical cutoffs for the numbers of proteins $p53$ and $A$: $p53_{\max} = C_{\text{p53}}$ for $p53(t)$ and $A_{\max} = C_{\text{A}}$ for $A(t)$. Without loss of generality, assume that $C_{\text{p53}} = C_{\text{A}} = C$ with $C$ being a known positive integer. In addition, we introduce an operator that will be used to the calculation of matrix $\mathbf{M}$ defined later.

In order to give the expression of $\mathbf{M}$, we also introduce operators, denoted by $\mathbb{L}^{(i)}$, $(i = 1,2,\cdots,n)$, which acts on matrices with the operation rule being as follows: $\mathbb{L}^{(i)}\mathbf{M} = \mathbf{M}^{(i)}$, where $\mathbf{M}^{(i)}$ is a matrix whose order is the same as $\mathbf{M}$ but some components are zero, e.g., if $\mathbf{M} = (a_{ij})_{3\times3}$, we obtain $\mathbb{L}^{(2)}\mathbf{M} = \mathbf{M}^{(2)} = (b_{ij})_{3\times3}$, where $(b_{ij})_{2\times2} = (a_{ij})_{2\times2}$ and the other elements are equal to zero. Specially, we always keep $\mathbb{L}^{(0)}\mathbf{M} = \mathbf{M}^{(0)} = \mathbf{0}$ if $i = 0$, where $\mathbf{0}$ has same order with $\mathbf{M}$.

Owing to big differences in analysis and calculation, we will separately discuss the cases of four kinds of absorbing domains.

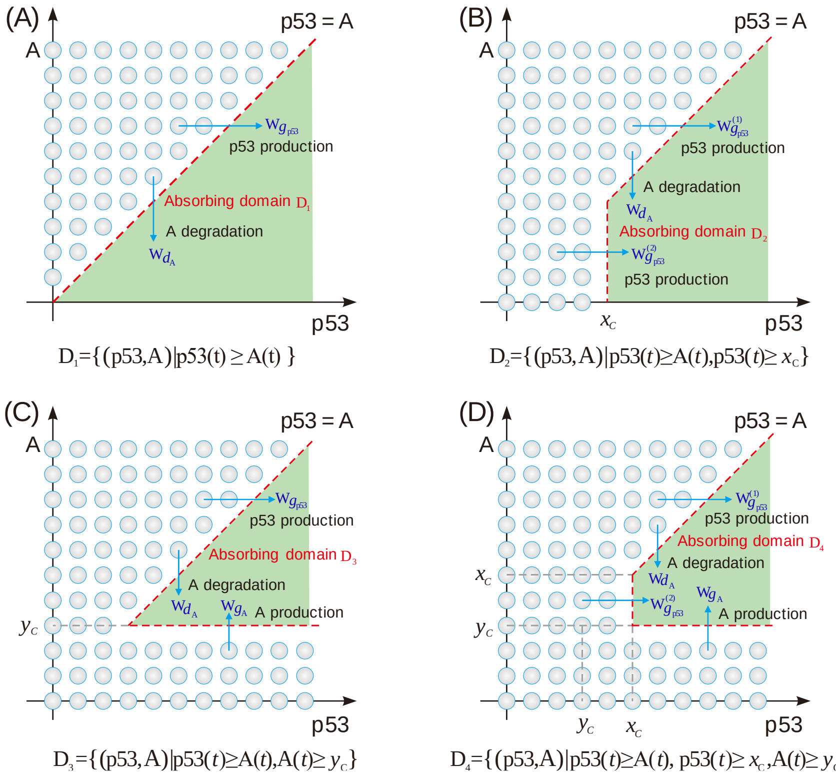

**Fig. S3 Schematic diagram for the possible patterns of the absorbing domain of variable** $p53$ **to reach barrier** $A$: (A) $\mathcal{D}_1 = \{(p53, A) | \, p53(t) \ge A(t)\}$; (B) $\mathcal{D}_2 = \{(p53, A) | \, p53(t) \ge A(t), p53(t) \ge x_C\}$; (C) $\mathcal{D}_3 = \{(p53, A) | \, p53(t) \ge A(t), A(t) \ge y_C\}$ (D) $\mathcal{D}_4 = \{(p53, A) | \, p53(t) \ge A(t), p53(t) \ge x_C, A(t) \ge y_C\}$. In figure, circles represent points in the state space, lines represent the boundary of absorbing domain, and arrows represent the direction of threshold crossing.

#### 3.1 The absorbing domain is $\mathcal{D}_1$

If the trajectory $\{(p53, A)\}$ of protein pair $p53$ and $A$ is considered in the domain $\{(p53, A) | \, p53 = p53(t) < A(t) = A\}$ (referring to Fig. S3A), the random variable $T$ by the time at which protein counts $p53$ reaches protein counts $A$ for the first time, is defined as

$$T = \min\{t : p53(t) \ge A(t) | \, p53(0) = m < A(0) = n\}. \tag{S28}$$

The absorbing domain in this case is given by

$$\mathcal{D}_1 = \left\{ (p53, A) \middle| p53(t) \geq A(t) \right\} \tag{S29}$$

And the corresponding finite state-space for birth-death process is defined as

$$\Omega = \left\{ (p53(t), A(t)) \middle| p53(t) < A(t), p53(t) = 0,1,2,\cdots,C-1, A(t) = 1,2,\cdots,C \right\}$$

If we write $P_{m,n}(t) \equiv P(p53(t) = m, A(t) = n)$, then its finite state-space is

$$\Omega = \left\{ (m,n) \middle| m < n, m = 0,1,2,\cdots,C-1, n = 1,2,\cdots,C \right\} \tag{S30}$$

The chemical master equation for $P_{m,n}(t)$ can thus be written in the form of Eq. S25 with $(m,n) \in \Omega$. For convenience, we introduce the denotation

$$\mathbf{P}_{m,n}(t) = \left[ \mathbf{P}_{*,1}, \mathbf{P}_{*,2}, \mathbf{P}_{*,3}, \cdots, \mathbf{P}_{*,C} \right]^{\mathrm{T}} = \left[ [P_{01}, 0, \cdots, 0], [P_{02}, P_{12}, 0, \cdots, 0], \cdots, [P_{0,C}, P_{1,C}, \cdots, P_{C-1,C}] \right]^{\mathrm{T}}$$

where the denotation $\mathbf{P}_{*,k} = \left[ P_{0,k}, P_{1,k}, \cdots, P_{k-1,k}, 0, \cdots, 0 \right]$, $k \in 1,2,\cdots,\mathrm{C}$.

Based on the sub-matrix operator $\mathbb{L}$, we can easily determine the matrix $\mathbf{M}$ in the master equation $\partial_t \mathbf{P}_{m,n}(t) = \mathbf{M}\mathbf{P}_{m,n}(t)$ or in Eq. S25 with $(m,n) \in \Omega$. The form of the matrix $\mathbf{M}$ is

$$\mathbf{M} = \begin{bmatrix} \mathbf{D}_1 & \mathbf{U}_1 & & & & \\ \mathbf{L}_1 & \mathbf{D}_2 & \mathbf{U}_2 & & & \\ & \mathbf{L}_2 & \mathbf{D}_3 & \mathbf{U}_3 & & \\ & & \ddots & \ddots & \ddots & \\ & & & \mathbf{L}_{C-2} & \mathbf{D}_{C-1} & \mathbf{U}_{C-1} \\ & & & & \mathbf{L}_{C-1} & \mathbf{D}_C \end{bmatrix} \tag{S31}$$

where

$$\begin{cases} \mathbf{U}_i = (i+1)\mathbb{L}^{(i)}(d_{\mathrm{A}}\mathbf{I}_C), & i = 1,2,\cdots,C-1. \\ \mathbf{D}_i = \mathbb{L}^{(i)}\left[ \mathbf{M}_{g_{\mathrm{p53}}} + \mathbf{M}_{d_{\mathrm{p53}}} - (g_{\mathrm{A}} + i d_{\mathrm{A}})\mathbf{I}_C \right], & i = 1,2,\cdots,C. \\ \mathbf{L}_i = \mathbb{L}^{(i)}(g_{\mathrm{A}}\mathbf{I}_C), & i = 1,2,\cdots,C-1. \end{cases}$$

Here $\mathbf{I}_C$ is the $C \times C$ identity matrix. And the matrix $\mathbf{M}_{d_{\mathrm{p53}}}$ is $\mathbf{M}_{d_{\mathrm{p53}}} = d_{\mathrm{p53}}\mathrm{diag}([1,2,\ldots,C-1],1) - d_{\mathrm{p53}}\mathrm{diag}([0,1\ldots,C-1],0)$ , where symbol $\mathrm{diag}(\mathbf{v},k)$ represents that the elements of vector $\mathbf{v}$ are placed on the $k$th diagonal, where $k = 0$ corresponds to the main diagonal, $k > 0$ to the upper principal diagonal, and $k < 0$ to the lower principal diagonal. That is,

$$\mathbf{M}_{d_{\mathrm{p53}}}=\begin{bmatrix}0 & d_{\mathrm{p53}} & & & & \\ & -d_{\mathrm{p53}} & 2d_{\mathrm{p53}} & & & \\ & & -2d_{\mathrm{p53}} & 3d_{\mathrm{p53}} & & \\ & & & \ddots & \ddots & \\ & & & & -(C-2)d_{\mathrm{p53}} & (C-1)d_{\mathrm{p53}} \\ & & & & & -(C-1)d_{\mathrm{p53}}\end{bmatrix}. \tag{S32}$$

Moreover, we have $\mathbf{M}_{g_{\mathrm{p53}}}=g_{\mathrm{p53}}\mathbf{M}_{\mathrm{burst}}$, where $\mathbf{M}_{\mathrm{burst}}=-P_{B\geq 1}\mathbf{I}_C+\sum_{k=1}^{C-1}P_{B=k}\mathrm{diag}\left(\mathbf{e}_{C-k}^{\mathrm{T}},-k\right)$. In the case of feedback, implying that $g_{\mathrm{p53}}$ depends on the molecule number ($m$) of protein $p53$, $\mathbf{M}_{g_{\mathrm{p53}}^{(m)}}=\mathbf{M}_{\mathrm{burst}}\mathbf{G}$, where $\mathbf{G}=\mathrm{diag}\left(\left[g_{\mathrm{p53}}^{(0)},g_{\mathrm{p53}}^{(1)}\cdots,g_{\mathrm{p53}}^{(C-1)}\right],0\right)$.

$$\mathbf{M}_{g_{\mathrm{p53}}}=\begin{bmatrix}-g_{\mathrm{p53}}^{(0)}P_{B\geq 1} & & & & & \\ g_{\mathrm{p53}}^{(0)}P_{B=1} & -g_{\mathrm{p53}}^{(1)}P_{B\geq 1} & & & & \\ g_{\mathrm{p53}}^{(0)}P_{B=2} & g_{\mathrm{p53}}^{(1)}P_{B=1} & -g_{\mathrm{p53}}^{(2)}P_{B\geq 1} & & & \\ \vdots & \vdots & \vdots & \ddots & & \\ g_{\mathrm{p53}}^{(0)}P_{B=C-2} & g_{\mathrm{p53}}^{(1)}P_{B=C-3} & g_{\mathrm{p53}}^{(2)}P_{B=C-4} & \cdots & -g_{\mathrm{p53}}^{(C-2)}P_{B\geq 1} & \\ g_{\mathrm{p53}}^{(0)}P_{B=C-1} & g_{\mathrm{p53}}^{(1)}P_{B=C-2} & g_{\mathrm{p53}}^{(2)}P_{B=C-3} & \cdots & g_{\mathrm{p53}}^{(C-2)}P_{B=1} & -g_{\mathrm{p53}}^{(C-1)}P_{B\geq 1}\end{bmatrix} \tag{S33}$$

Given a numerical cutoff $(C)$, the FPT distribution can thus be determined by the following way. Apparently, the probability of a state reaches the absorption domain given by $\mathcal{D}_1=\left\{(p53,A)\middle| p53(t)\geq A(t)\right\}$ in the infinitesimal time interval $(t,t+dt)$, is the sum of the following two terms: the first one is the probability that $\left\{p53(t)=m,A(t)=n\right\}$ and a jump of size $n-m$ or large occurs in the time interval $[t,t+dt)$, and the second one is the degradation probability that $\left\{p53(t)=n-1,A(t)=n\right\}$ occurs in the time interval $[t,t+dt)$. Thus, the probability density function of the first passage time (FPT) that a pair of proteins $(p53,A)$ reach absorbing domain $\mathcal{D}_1$ is given by

$$\begin{aligned} f_T(t) &= \sum_{m=0}^{C-1}(m+1)d_{\mathrm{A}}P_{m,m+1}(t)+\sum_{n=1}^{C}\sum_{m=0}^{n-1}g_{\mathrm{p53}}^{(m)}P_{B\geq n-m}P_{m,n}(t) \\ &= \sum_{\substack{n=1\\ m=n-1}}^{C} nd_{\mathrm{A}}P_{m,n}(t)+\sum_{n=1}^{C}\sum_{m=0}^{n-1}g_{\mathrm{p53}}^{(m)}P_{B\geq n-m}P_{m,n}(t) \\ &= \mathbf{W}_{d_{\mathrm{A}}}^{\mathrm{T}}\mathbf{P}_{m,n}(t)+\mathbf{W}_{g_{\mathrm{p53}}}^{\mathrm{T}}\mathbf{P}_{m,n}(t) \\ &\equiv \mathbf{W}^{\mathrm{T}}\mathbf{P}_{m,n}(t) \end{aligned} \tag{S34}$$

where we denote the column vector $\mathbf{W}_{d_{\mathrm{A}}}=\left(\mathbf{W}_{d_{\mathrm{y}}}^{n}\right)_{C\times 1}$ with $\mathbf{W}_{d_{\mathrm{A}}}^{n}=nd_{\mathrm{A}}\mathbf{1}_n$, $n=1,2\cdots,C$, and define a column vector of length $C$, $\mathbf{1}_i = \left(0,\cdots 0,1,0,\cdots,0\right)^{\mathrm{T}}$, in which the only $i$ th element is equal to 1 and other elements are all zero. Similarly, the column vector $\mathbf{W}_{g_{\mathrm{p53}}}$ can be rewritten as $\mathbf{W}_{g_{\mathrm{p53}}} = \left(\mathbf{W}^n_{g_{\mathrm{p53}}}\right)_{C\times 1}$ with $\mathbf{W}^n_{g_{\mathrm{p53}}} = \sum_{m=1}^{n-1} g_{\mathrm{p53}}^{(m)} P_{B\geq n-m} \mathbf{1}_{m+1}$, $n = 1,2\cdots,C$. Thus, the column vector $\mathbf{W}$ can be expressed as

$$\mathbf{W} = \mathbf{W}_{g_{\mathrm{p53}}} + \mathbf{W}_{d_{\mathrm{A}}} = \left[\mathbf{W}_1^{\mathrm{T}}, \mathbf{W}_2^{\mathrm{T}}, \cdots, \mathbf{W}_C^{\mathrm{T}}\right]^{\mathrm{T}} \tag{S35}$$

where $\mathbf{W}_n = \mathbf{W}^n_{d_{\mathrm{A}}} + \mathbf{W}^n_{g_{\mathrm{p53}}} = n d_{\mathrm{A}} \mathbf{1}_n + \sum_{m=0}^{n-1} g_{\mathrm{p53}}^{(m)} P_{B\geq n-m} \mathbf{1}_{m+1}$, $n = 1,2\cdots,C$. In brevity, the involved vectors can be expressed as

$$\mathbf{W}^{\mathrm{T}}_{d_{\mathrm{A}}} = \left[\left[d_{\mathrm{A}},0,\cdots,0\right],\left[0,2d_{\mathrm{A}},0,\cdots,0\right],\cdots,\left[0,0,\cdots,0,Cd_{\mathrm{A}}\right]\right],$$

$$\mathbf{W}^{\mathrm{T}}_{g_{\mathrm{p53}}} = \left[\left[g_{\mathrm{p53}}^{(0)} P_{B\geq 1},0,\cdots,0\right],\left[g_{\mathrm{p53}}^{(0)} P_{B\geq 2}, g_{\mathrm{p53}}^{(1)} P_{B\geq 1},0,\cdots,0\right],\cdots,\left[g_{\mathrm{p53}}^{(0)} P_{B\geq C}, g_{\mathrm{p53}}^{(1)} P_{B\geq C-1},\cdots, g_{\mathrm{p53}}^{(C-1)} P_{B\geq 1}\right]\right],$$

$$\mathbf{W}^{\mathrm{T}} = \left[\left[g_{\mathrm{p53}}^{(0)} P_{B\geq 1} + d_{\mathrm{A}},0,\cdots,0\right],\left[g_{\mathrm{p53}}^{(0)} P_{B\geq 2}, g_{\mathrm{p53}}^{(1)} P_{B\geq 1} + 2d_{\mathrm{A}},0,\cdots,0\right],\cdots,\left[g_{\mathrm{p53}}^{(0)} P_{B\geq C}, g_{\mathrm{p53}}^{(1)} P_{B\geq C-1},\cdots, g_{\mathrm{p53}}^{(C-1)} P_{B\geq 1} + Cd_{\mathrm{A}}\right]\right]$$

The formulation for calculating moments of the FPT is the same as Eq. S17.

Numerical results for mean FPT and timing variability are shown in Figs. S4 and S5.

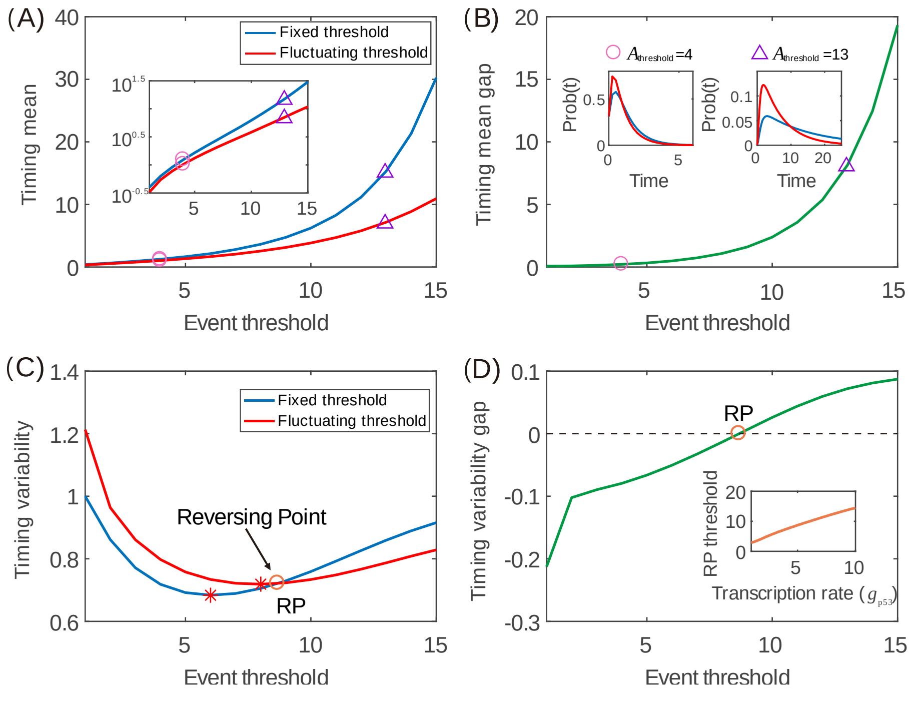

**Fig. S4 Comparison between the effects of fixed and fluctuating thresholds on timing**. (A) Timing mean as a function of event threshold for two different kinds of thresholds, where the inset shows timing mean as a function of event threshold on the logarithmic scale. (B) A different demonstration of the results in (A), showing the difference of the timing mean in the case of fixed threshold minus that in the case of fluctuating threshold, where insets show FPT distributions for two different event thresholds (indicated by empty circle and triangle) corresponding to $A_{\text{thresold}} = 4$ and $A_{\text{thresold}} = 13$. (C) Timing variability as a function of event threshold for two different kinds of thresholds, where the empty circle is the crossing point of two curves, and stars represent the critical threshold that makes timing variability reach the minimum. (D) As a supplement of (C), the difference of timing variability in the case of fixed threshold minus that in the case of fluctuating threshold, where the inset shows the critical threshold as a function of the transcription rate. In (A) and (C), the parameter values are set as $g_{\text{p53}} = 5$, $d_{\text{p53}} = 1$, $b = 1$, $p53_{\text{max}} = 30$, the fixed threshold is $A_{\text{threshold}} = 10$, and the fluctuating threshold corresponds to $g_{\text{A}} = 10$ and $d_{\text{A}} = 1$. The inset in (D) corresponds to $d_{\text{p53}} = 1$, $b = 1$, $g_{\text{A}} = 10$, $d_{\text{A}} = 1$, $g_{\text{p53}} = 1 \sim 10$ and $p53_{\text{max}} = 30$.

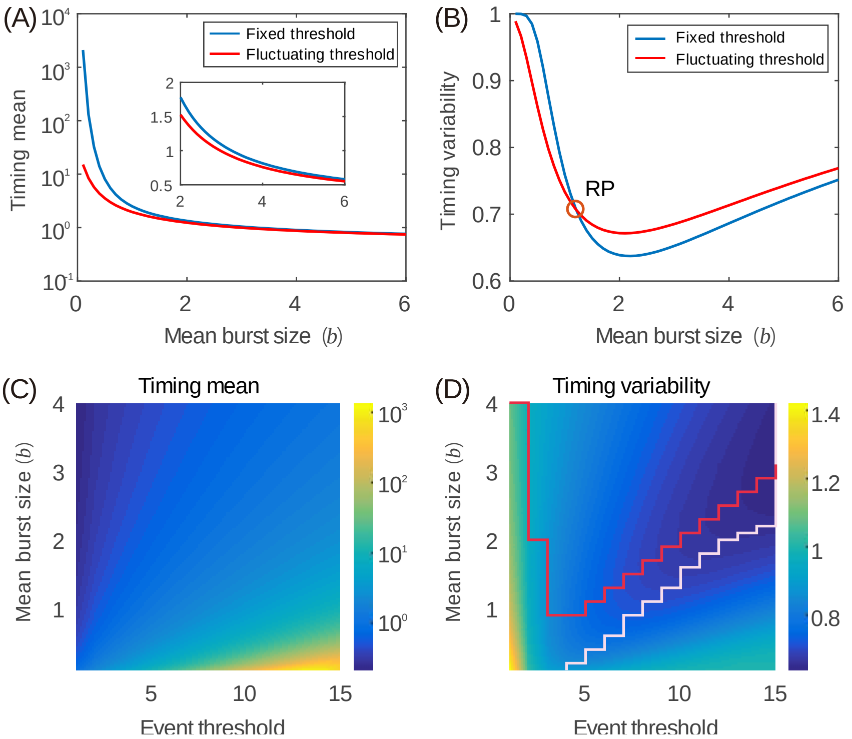

**Fig. S5 Effect of mean burst sizes on event timing**. (A) Timing mean as a function of the mean burst size, where the inset shows a partial enlarged diagram. (B) Timing variability as a function of the mean burst size two different cases: fixed threshold (blue curve) and fluctuating threshold (red curve), where RP represents a critical point for reversing. (C) Heatmap showing timing mean as a function of event threshold and mean burst size. (D) Heatmap showing timing variability as a function of event threshold and mean burst size, where the red curve corresponds to the case that an event of threshold crossing has happened either due to a sufficiently large transcription rate ($g_{\text{p53}}$) or due to a sufficiently small degradation rate ($d_{\text{p53}}$), and the stair-like line consists of the points corresponding to the star in Fig. 3*C* in the main file wherein a special value of $g_{\text{p53}}$ is used. In (A)-(D), the parameter values are set as $g_{\text{p53}} = 5$, $d_{\text{p53}} = 1$, $p53_{\text{max}} = 30$. In (A) and (B), the threshold is set as $A_{\text{thresold}} = g_{\text{A}} / d_{\text{A}}$, with $g_{\text{A}} = 10$ and $d_{\text{A}} = 1$. whereas in (C) and (D), $d_{\text{A}} = 1$ and $g_{\text{A}}$ is changed in $A_{\text{thresold}} = g_{\text{A}} / d_{\text{A}}$.

##### 3.2 The absorbing domain is $\mathcal{D}_2$

If the trajectory $(p53, A)$ of protein pair $p53$ and $A$ is considered in two domains $\{(p53, A) | p53(t) < A(t)\}$ and $\{(p53, A) | p53(t) < x_C\}$ (referring to Fig. S3*B*), we define the random variable $T$ by the time at which protein counts $p53$ reach protein counts $A$ for the first time, that is,

$$T = \min\{t : p53(t) \geq A(t), p53(t) \geq x_C \mid p53(0) = m < A(0) = n\} \tag{S36}$$

The absorbing domain in this case is given by

$$\mathcal{D}_2 = \{(p53(t), A(t)) | p53(t) \geq A(t), p53(t) \geq x_C\}. \tag{S37}$$

The corresponding finite state-space for birth-death process $\Omega$ is defined as $\Omega = \Omega_1 \cup \Omega_2$, where

$$\Omega_1 = \{(p53(t), A(t)) | p53(t) < x_C, p53(t) = 0,1,2,\cdots,x_C - 1, A(t) = 0,1,2,\cdots,C\},$$

$$\Omega_2 = \{(p53(t), A(t)) | p53(t) < A(t), p53(t) = 0,1,2,\cdots,C-1, A(t) = 0,1,2,\cdots,C\}.$$

Then, we easily write the finite state-space $\Omega$, which consists of two parts $\Omega_1, \Omega_2$, i.e., $\Omega = \Omega_1 \cup \Omega_2$, where we denote respectively

$$\Omega_1 = \{(m,n) | m = 0,1,2,\cdots,x_C - 1, n = 0,1,2,\cdots,C\}, \tag{S38}$$

$$\Omega_2 = \{(m,n) | m < n, m = 0,1,2,\cdots,C-1, n = 1,2,\cdots,C\}. \tag{S39}$$

The corresponding chemical master equation for $P_{m,n}(t)$ satisfies the form of Eq. S25 with $(m,n) \in \Omega$. Denote

$$\mathbf{P}_{m,n}(t)=\left[\mathbf{P}_{*,0},\mathbf{P}_{*,1},\cdots,\mathbf{P}_{*,x_C-1},\mathbf{P}_{*,x_C},\mathbf{P}_{*,x_C+1},\cdots,\mathbf{P}_{*,C}\right]^{\mathrm{T}}$$
$$=\Big[\left[P_{0,0},P_{1,0},\cdots,P_{x_C-1,0},0,\cdots,0\right],\cdots,\left[P_{0,x_C-1},P_{1,x_C-1},\cdots,P_{x_C-1,x_C-1},0,\cdots,0\right],$$
$$\left[P_{0,x_C},P_{1,x_C},\cdots,P_{x_C-1,x_C},0,\cdots,0\right],\left[P_{0,x_C+1},\cdots,P_{x_C-1,x_C+1},P_{x_C,x_C+1},0,\cdots,0\right],$$
$$\cdots,\left[P_{0,C},P_{1,C},\cdots,P_{C-1,C}\right]\Big]^{\mathrm{T}}$$

Based on the sub-matrix operator $\mathbb{L}$, we can easily determine matrix $\mathbf{M}$ in the master equation $\partial_t\mathbf{P}_{m,n}(t)=\mathbf{M}\mathbf{P}_{m,n}(t)$ or in Eq. S25 with $(m,n)\in\Omega$. The form of the matrix $\mathbf{M}$ is

$$\mathbf{M}=\begin{bmatrix}\mathbf{D}_0 & \mathbf{U}_0 & & & & & \\ \mathbf{L}_0 & \mathbf{D}_1 & \mathbf{U}_1 & & & & \\ & \mathbf{L}_1 & \mathbf{D}_2 & \mathbf{U}_2 & & & \\ & & \ddots & \ddots & \ddots & & \\ & & & \mathbf{L}_{C-2} & \mathbf{D}_{C-1} & \mathbf{U}_{C-1} \\ & & & & \mathbf{L}_{C-1} & \mathbf{D}_C\end{bmatrix} \tag{S40}$$

where,

$$\mathbf{U}_i\equiv\begin{cases}(i+1)\left(\mathbb{L}^{(x_C)}\left(d_{\mathrm{A}}\mathbf{I}_C\right)\right), & i=0,1,\cdots,x_C-1;\\ (i+1)\left(\mathbb{L}^{(i)}\left(d_{\mathrm{A}}\mathbf{I}_C\right)\right), & i=x_C,x_C+1,\cdots,C-1.\end{cases}$$

$$\mathbf{D}_i\equiv\begin{cases}\mathbb{L}^{(x_C)}\left(\mathbf{M}_{g_{\mathrm{p53}}}+\mathbf{M}_{d_{\mathrm{p53}}}-\left(g_{\mathrm{A}}+id_{\mathrm{A}}\right)\mathbf{I}_C\right), & i=0,1,\cdots,x_C-1;\\ \mathbb{L}^{(i)}\left(\mathbf{M}_{g_{\mathrm{p53}}}+\mathbf{M}_{d_{\mathrm{p53}}}-\left(g_{\mathrm{A}}+id_{\mathrm{A}}\right)\mathbf{I}_C\right), & i=x_C,x_C+1,\cdots,C\end{cases}$$

$$\mathbf{L}_i\equiv\begin{cases}\mathbb{L}^{(x_C)}\left(g_{\mathrm{A}}\mathbf{I}_C\right), & i=0,1,\cdots,x_C-1;\\ \mathbb{L}^{(i)}\left(g_{\mathrm{A}}\mathbf{I}_C\right), & i=x_C,x_C+1,\cdots,C-1.\end{cases}$$

Here, $\mathbf{I}_C$, $\mathbf{M}_{d_{\mathrm{p53}}}$ and $\mathbf{M}_{g_{\mathrm{p53}}}$ have the same definition as the case with absorbing domain $\mathcal{D}_1$.

Note that the probability that a state reaches the absorbing domain defined by $\mathcal{D}_2=\left\{\left(p53(t),A(t)\right)\middle|\,p53(t)\geq A(t),p53(t)\geq x_C\right\}$ in the infinitesimal time interval $(t,t+dt)$ is the sum of the following three terms: the probability that $\left\{p53(t)=m,A(t)=n\right\}$ and a jump of size $n-m$ or large occurs in the time interval $[t,t+dt)$, the degradation probability that $\left\{p53(t)=n-1,A(t)=n\right\}$ occurs in the time interval $[t,t+dt)$, and the probability that $\left\{p53(t)=m,A(t)=n\right\}$ and a jump of size $x_C-m$ or large occurs in the time interval $[t,t+dt)$. Thus, the probability density function of the FPT that $\left(p53(t),A(t)\right)$ reaches absorbing domain $\mathcal{D}_2$ is given by

$$
\begin{aligned}
f_T(t) &= \sum_{m=x_C}^{C-1}(m+1)d_{\mathrm{A}}P_{m,m+1}(t)+\sum_{n=x_C}^{C}\sum_{m=0}^{n-1}g_{\mathrm{p53}}^{(m)}P_{B\geq n-m}P_{m,n}(t)+\sum_{n=0}^{x_C-1}\sum_{m=0}^{x_C-1}g_{\mathrm{p53}}^{(m)}P_{B\geq x_C-m}P_{m,n}(t)\\
&= \sum_{\substack{n=x_C+1\\ m=n-1}}^{C} nd_{\mathrm{A}}P_{m,n}(t)+\sum_{n=x_C}^{C}\sum_{m=0}^{n-1}g_{\mathrm{p53}}^{(m)}P_{B\geq n-m}P_{m,n}(t)+\sum_{n=0}^{x_C-1}\sum_{m=0}^{x_C-1}g_{\mathrm{p53}}^{(m)}P_{B\geq x_C-m}P_{m,n}(t)\\
&= \mathbf{W}_{d_{\mathrm{A}}}^{\mathrm{T}}\mathbf{P}_{m,n}(t)+\mathbf{W}_{g_{\mathrm{p53}}}^{(1)\mathrm{T}}\mathbf{P}_{m,n}(t)+\mathbf{W}_{g_{\mathrm{p53}}}^{(2)\mathrm{T}}\mathbf{P}_{m,n}(t)\\
&\equiv \mathbf{W}^{\mathrm{T}}\mathbf{P}_{m,n}(t)
\end{aligned}
\tag{S41}
$$

where we denote respectively three column vectors

$$
\begin{cases}
\mathbf{W}_{d_{\mathrm{A}}} = \left(\mathbf{W}_{d_{\mathrm{A}}}^{n}\right)_{(C+1)\times 1}\\
\mathbf{W}_{g_{\mathrm{p53}}}^{(1)} = \left(\mathbf{W}_{g_{\mathrm{p53}}}^{n}\right)_{(C+1)\times 1}\\
\mathbf{W}_{g_{\mathrm{p53}}}^{(2)} = \left(\hat{\mathbf{W}}_{g_{\mathrm{p53}}}^{n}\right)_{(C+1)\times 1}
\end{cases}
$$

which the corresponding vector is given by

$$
\mathbf{W}_{d_{\mathrm{A}}}^{n} = \begin{cases}\mathbf{0}, & n=0,1,\cdots,x_C;\\ nd_{\mathrm{A}}\mathbf{1}_n, & n=x_C+1,x_C+2,\cdots,C.\end{cases}
$$

$$
\mathbf{W}_{g_{\mathrm{p53}}}^{n} = \begin{cases}\mathbf{0}, & n=0,1,\cdots,x_C-1;\\ \sum_{m=0}^{n-1}g_{\mathrm{p53}}^{(m)}P_{B\geq n-m}\mathbf{1}_{m+1}, & n=x_C,x_C+1,\cdots,C.\end{cases}
$$

$$
\hat{\mathbf{W}}_{g_{\mathrm{p53}}}^{n} = \begin{cases}\sum_{m=0}^{x_C-1}g_{\mathrm{p53}}^{(m)}P_{B\geq x_C-m}\mathbf{1}_{m+1}, & n=0,1,\cdots,x_C-1;\\ \mathbf{0}, & n=x_C,x_C+1,\cdots,C.\end{cases}
$$

Thus, the column vector $\mathbf{W}$ takes the following form

$$
\mathbf{W}=\mathbf{W}_{g_{\mathrm{p53}}}^{(1)}+\mathbf{W}_{g_{\mathrm{p53}}}^{(2)}+\mathbf{W}_{d_{\mathrm{A}}}=\left[\mathbf{W}_0^{\mathrm{T}},\mathbf{W}_1^{\mathrm{T}},\mathbf{W}_2^{\mathrm{T}},\cdots,\mathbf{W}_n^{\mathrm{T}}\right]^{\mathrm{T}}
\tag{S42}
$$

where $\mathbf{W}_n=\mathbf{W}_{g_{\mathrm{p53}}}^{n}+\hat{\mathbf{W}}_{g_{\mathrm{p53}}}^{n}+\mathbf{W}_{d_{\mathrm{A}}}^{n}$, which can be expressed as

$$
\mathbf{W}_n = \begin{cases}\sum_{m=0}^{x_C-1}g_{\mathrm{p53}}^{(m)}P_{B\geq x_C-m}\mathbf{1}_{m+1}, & n=0,1,\cdots,x_C-1,x_C;\\ nd_{\mathrm{A}}\mathbf{1}_n+\sum_{m=0}^{n-1}g_{\mathrm{p53}}^{(m)}P_{B\geq n-m}\mathbf{1}_{m+1}, & n=x_C+1,x_C+2\cdots,C.\end{cases}
$$

In brevity, the involved vectors can be expressed as

$$
\mathbf{W}_{d_{\mathrm{A}}}^{\mathrm{T}}=\left[[0,\cdots,0],\cdots,[0,\cdots,0],\left[0,\cdots,0,(x_C+1)d_{\mathrm{A}},0,\cdots,0\right],\left[0,\cdots,0,(x_C+2)d_{\mathrm{A}},0,\cdots,0\right],\cdots,\left[0,\cdots,0,Cd_{\mathrm{A}}\right]\right]
$$

$$
\begin{aligned}
\mathbf{W}_{g_{\mathrm{p53}}}^{(1)\mathrm{T}}=\Big[&[0,\cdots,0],\cdots,[0,\cdots,0],\left[g_{\mathrm{p53}}^{(0)}P_{B\geq x_C},g_{\mathrm{p53}}^{(1)}P_{B\geq x_C-1},\cdots,g_{\mathrm{p53}}^{(x_C-1)}P_{B\geq 1},0,\cdots,0\right],\\
&\left[g_{\mathrm{p53}}^{(0)}P_{B\geq x_C+1},g_{\mathrm{p53}}^{(1)}P_{B\geq x_C},\cdots,g_{\mathrm{p53}}^{(x_C)}P_{B\geq 1},0,\cdots,0\right],\cdots,\left[g_{\mathrm{p53}}^{(0)}P_{B\geq C},g_{\mathrm{p53}}^{(1)}P_{B\geq C-1},\cdots,g_{\mathrm{p53}}^{(C-1)}P_{B\geq 1}\right]\Big]
\end{aligned}
$$

$$
\begin{aligned}
\mathbf{W}_{g_{\mathrm{p53}}}^{(2)\mathrm{T}}=\Big[&\left[g_{\mathrm{p53}}^{(0)}P_{B\geq x_C},g_{\mathrm{p53}}^{(1)}P_{B\geq x_C-1},\cdots,g_{\mathrm{p53}}^{(x_C-1)}P_{B\geq 1},0,\cdots,0\right],\left[g_{\mathrm{p53}}^{(0)}P_{B\geq x_C},g_{\mathrm{p53}}^{(1)}P_{B\geq x_C-1},\cdots,g_{\mathrm{p53}}^{(x_C-1)}P_{B\geq 1},0,\cdots,0\right],\cdots,\\
&\left[g_{\mathrm{p53}}^{(0)}P_{B\geq x_C},g_{\mathrm{p53}}^{(1)}P_{B\geq x_C-1},\cdots,g_{\mathrm{p53}}^{(x_C-1)}P_{B\geq 1},0,\cdots,0\right],[0,\cdots,0],\cdots,[0,\cdots,0]\Big]
\end{aligned}
$$

$$\mathbf{W}^{\mathrm{T}} = \Big[\Big[ g_{\mathrm{p53}}^{(0)} P_{B\geq x_C}, g_{\mathrm{p53}}^{(1)} P_{B\geq x_C-1}, \cdots, g_{\mathrm{p53}}^{(x_C-1)} P_{B\geq 1}, 0, \cdots, 0 \Big], \cdots, \Big[ g_{\mathrm{p53}}^{(0)} P_{B\geq x_C}, g_{\mathrm{p53}}^{(1)} P_{B\geq x_C-1}, \cdots, g_{\mathrm{p53}}^{(x_C-1)} P_{B\geq 1}, 0, \cdots, 0 \Big],$$
$$\Big[ g_{\mathrm{p53}}^{(0)} P_{B\geq x_C+1}, g_{\mathrm{p53}}^{(1)} P_{B\geq x_C}, \cdots, (x_C+1) d_{\mathrm{A}} + g_{\mathrm{p53}}^{(x_C)} P_{B\geq 1}, 0, \cdots, 0 \Big], \cdots, \Big[ g_{\mathrm{p53}}^{(0)} P_{B\geq C}, g_{\mathrm{p53}}^{(1)} P_{B\geq C-1}, \cdots, C d_{\mathrm{A}} + g_{\mathrm{p53}}^{(C-1)} P_{B\geq 1} \Big]\Big].$$

The formulation for calculating the moments of the FPT is the same as Eq. S17.

Numerical results for mean FPT and timing variability are shown in Fig. S6.

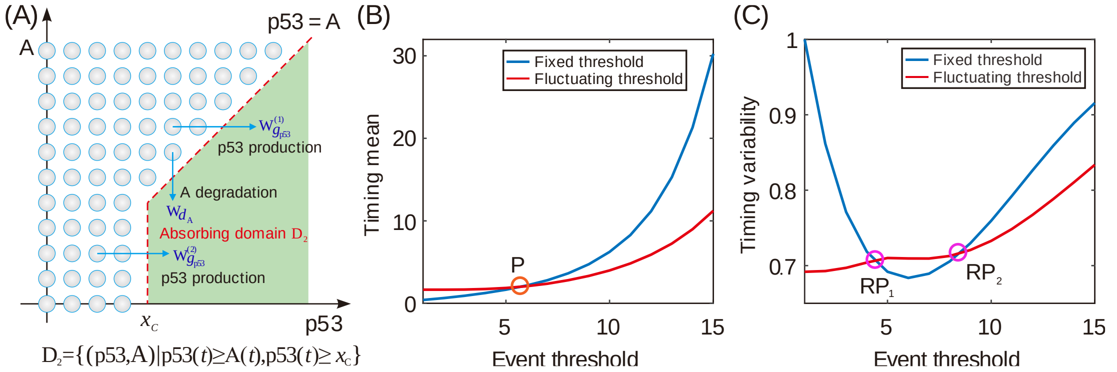

**Fig.S6 Characteristic of the curve for timing mean or timing variability vs event threshold in the case of absorbing domain** $\mathcal{D}_2$. (A) Schematic for PFT when an absorbing domain is defined by $\mathcal{D}_2 = \{(p53, A) | \, p53(t) \geq A(t), p53(t) \geq x_C\}$, where arrows represent the direction of threshold crossing. (B) Timing mean as a function of event threshold for two different kinds of thresholds, where empty circle represents the crossing point $\mathrm{P}$ of two curves for the mean FPT in two cases of event threshold. (C) Timing variability as a function of event threshold for two different kinds of thresholds, where two empty circles are the crossing points of two curves, denoted by $\mathrm{RP}_1$ and $\mathrm{RP}_2$. In (B) and (C), the parameter values are set as $x_{\mathrm{C}} = 5$, $g_{\mathrm{p53}} = 5$, $d_{\mathrm{p53}} = 1$, $b = 1$, the cutoff constant is set as $p53_{\max} = 50$. If the fixed threshold is set as $A_{\mathrm{threshold}} = 10$, the fluctuating threshold corresponds to $g_{\mathrm{A}} = 10$ and $d_{\mathrm{A}} = 1$.

#### 3.3 The absorbing domain is $\mathcal{D}_3$

If the trajectory $(p53, A)$ of protein pair $p53$ and $A$ is considered in two domain $\{(p53, A) | \, p53(t) < A(t), A(t) \geq y_C\}$ and $\{(p53, A) | \, A(t) < y_C\}$ (referring to Fig. S3C), we define the random variable $T$ as the time that protein $p53$ counts reach protein $A$ counts for the first time, that is,

$$T = \min\left\{t : p53(t) \geq A(t), A(t) \geq y_C \middle| p53(0) = m < A(0) = n\right\} \tag{S43}$$

The absorbing domain in this case is given by

$$\mathcal{D}_3 = \left\{(p53, A) \middle| p53(t) \geq A(t), A(t) \geq y_C\right\}. \tag{S44}$$

The corresponding finite state-space for birth-death process $\Omega$ with $\Omega = \Omega_1 \cup \Omega_2$, where

$$\Omega_1 = \left\{(p53, A) \middle| A(t) < y_C, p53(t) = 0,1,2,\cdots,C-1, A(t) = 0,1,2,\cdots,y_C - 1\right\},$$

$$\Omega_2 = \left\{(p53, A) \middle| p53(t) < Y(t), A(t) \geq y_C, p53(t) = 0,1,2,\cdots,C-1, A(t) = y_C, y_C + 1,\cdots,C\right\}.$$

Then we easily write its finite state-space $\Omega$, which consists of two parts $\Omega_1, \Omega_2$, i.e., $\Omega = \Omega_1 \cup \Omega_2$, where we denote respectively

$$\Omega_1 = \left\{(m,n) \middle| m = 0,1,2,\cdots,C-1, n = 0,1,2,\cdots,y_C - 1\right\}, \tag{S45}$$

$$\Omega_2 = \left\{(m,n) \middle| m < n, m = 0,1,2,\cdots,C-1, n = y_C, y_C + 1,\cdots,C\right\}. \tag{S46}$$

The corresponding chemical master equation for $P_{m,n}(t)$ takes the form of Eq.S25 with $(m,n) \in \Omega$.

Denote

$$\begin{aligned}\mathbf{P}_{m,n}(t) &= \left[\mathbf{P}_{*,0}, \mathbf{P}_{*,1}, \cdots, \mathbf{P}_{*,y_C-1}, \mathbf{P}_{*,y_C}, \mathbf{P}_{*,y_C+1}, \cdots, \mathbf{P}_{*,C}\right]^{\mathrm{T}} \\ &= \left[\left[P_{0,0}, P_{1,0}, \cdots, P_{C-1,0}\right], \cdots, \left[P_{0,y_C-1}, P_{1,y_C-1}, \cdots, P_{C-1,y_C-1}\right], \left[P_{0,y_C}, P_{1,y_C}, \cdots, P_{y_C-1,y_C}, 0, \cdots, 0\right],\right. \\ &\quad \left.\left[P_{0,y_C+1}, P_{1,y_C+1}, \cdots, P_{y_C,y_C+1}, 0, \cdots, 0\right], \cdots, \left[P_{0,C}, P_{1,C}, \cdots, P_{C-1,C}\right]\right]^{\mathrm{T}}\end{aligned}$$

Note that here $\mathbf{P}_{*,k}$ has two forms,

$$\begin{cases}\mathbf{P}_{*,\mathrm{k}} = \left[P_{0,k}, P_{1,k}, \cdots, P_{C-1,k}\right], & k = 0,1,\cdots,y_C - 1; \\ \mathbf{P}_{*,\mathrm{k}} = \left[P_{0,k}, P_{1,k}, \cdots, P_{k-1,k}, 0, \cdots, 0\right], & k = y_C, y_C + 1, \cdots, C.\end{cases}$$

It is obvious that the matrix $\mathbf{M}$, which satisfies the master equation $\partial_t \mathbf{P}_{m,n}(t) = \mathbf{M}\mathbf{P}_{m,n}(t)$ or in Eq. S25 with $(m,n) \in \Omega$, is also easily determined for the sub-matrix operator $\mathbb{L}$ in this case, i.e.,

$$\mathbf{M} = \begin{bmatrix} \mathbf{D}_0 & \mathbf{U}_0 & & & & \\ \mathbf{L}_0 & \mathbf{D}_1 & \mathbf{U}_1 & & & \\ & \mathbf{L}_1 & \mathbf{D}_2 & \mathbf{U}_2 & & \\ & & \ddots & \ddots & \ddots & \\ & & & \mathbf{L}_{C-2} & \mathbf{D}_{C-1} & \mathbf{U}_{C-1} \\ & & & & \mathbf{L}_{C-1} & \mathbf{D}_C \end{bmatrix} \tag{S47}$$

where,

$$
\mathbf{U}_i = \begin{cases} (i+1)\, d_{\mathrm{A}} \mathbf{I}_C, & i = 0,1,\cdots, y_C - 2; \\ (i+1)\left(\mathbb{L}^{(y_C)}\left(d_{\mathrm{A}} \mathbf{I}_C\right)\right), & i = y_C - 1; \\ (i+1)\left(\mathbb{L}^{(i)}\left(d_{\mathrm{A}} \mathbf{I}_C\right)\right), & i = y_C, y_C + 1, \cdots, C-1. \end{cases}
$$

$$
\mathbf{D}_i = \begin{cases} \mathbf{M}_{g_{\mathrm{p53}}} + \mathbf{M}_{d_{\mathrm{p53}}} - \left(g_{\mathrm{A}} + i d_{\mathrm{A}}\right)\mathbf{I}_C, & i = 0,1,\cdots, y_C - 1; \\ \mathbb{L}^{(i)}\left(\mathbf{M}_{g_{\mathrm{p53}}} + \mathbf{M}_{d_{\mathrm{p53}}} - \left(g_{\mathrm{A}} + i d_{\mathrm{A}}\right)\mathbf{I}_C\right), & i = y_C, y_C + 1, \cdots, C. \end{cases}
$$

$$
\mathbf{L}_i = \begin{cases} g_{\mathrm{A}} \mathbf{I}_C, & i = 0,1,\cdots, y_C - 2; \\ \mathbb{L}^{(y_C)}\left(g_{\mathrm{A}} \mathbf{I}_C\right), & i = y_C - 1; \\ \mathbb{L}^{(i)}\left(g_{\mathrm{A}} \mathbf{I}_C\right), & i = y_C, y_C + 1, \cdots, C-1. \end{cases}
$$

Here, $\mathbf{I}_C$, $\mathbf{M}_{d_{\mathrm{p53}}}$ and $\mathbf{M}_{g_{\mathrm{p53}}}$ have same definition with above case with absorbing domain $\mathcal{D}_1$.

Similarly, given a numerical cutoff $(C)$, we can determine the probability that a state reaches the absorbing domain defined by $\mathcal{D}_3 = \left\{(p53, A) \middle| p53(t) \geq A(t), A(t) \geq y_C\right\}$ in the infinitesimal time interval $(t, t+dt)$ in this case. This probability is the sum of the following three terms: the probability that $\left\{p53(t) = m, A(t) = n\right\}$ and a jump of size $n - m$ or larger occurs in the time interval $[t, t+dt)$, the degradation probability that $\left\{p53(t) = n-1, A(t) = n\right\}$ occurs in the time interval $[t, t+dt)$, and the transcription probability that $\left\{p53(t) = m, A(t) = y_C - 1\right\}$ with $m \geq y_C$ occurs in the time interval $[t, t+dt)$. Thus, the probability density function of the FPT that $(p53, A)$ reach absorbing domain $\mathcal{D}_3$ is given by

$$
\begin{aligned}
f_T(t) &= \sum_{\substack{n=y_C+1 \\ m=n-1}}^{C} n d_{\mathrm{A}} P_{m,n}(t) + \sum_{n=y_C}^{C} \sum_{m=0}^{n-1} g_{\mathrm{p53}}^{(m)} P_{B \geq n-m} P_{m,n}(t) + \sum_{\substack{m=y_C \\ n=y_C-1}}^{C-1} g_{\mathrm{A}} P_{m,n}(t) \\
&= \mathbf{W}_{d_{\mathrm{A}}}^{\mathrm{T}} \mathbf{P}_{m,n}(t) + \mathbf{W}_{g_{\mathrm{p53}}}^{\mathrm{T}} \mathbf{P}_{m,n}(t) + \mathbf{W}_{g_{\mathrm{A}}}^{\mathrm{T}} \mathbf{P}_{m,n}(t) \\
&\equiv \mathbf{W}^{\mathrm{T}} \mathbf{P}_{m,n}(t)
\end{aligned} \tag{S48}
$$

where we denote respectively three column vectors

$$
\begin{cases} \mathbf{W}_{d_{\mathrm{A}}} = \left(\mathbf{W}_{d_{\mathrm{A}}}^{n}\right)_{(C+1)\times 1} \\ \mathbf{W}_{g_{\mathrm{p53}}} = \left(\mathbf{W}_{g_{\mathrm{p53}}}^{n}\right)_{(C+1)\times 1} \\ \mathbf{W}_{g_{\mathrm{A}}} = \left(\mathbf{W}_{g_{\mathrm{A}}}^{n}\right)_{(C+1)\times 1} \end{cases}
$$

with

$$\mathbf{W}_{d_{\mathrm{A}}}^{n}=\begin{cases}\mathbf{0}, & n=0,1,\cdots,y_C;\\ nd_{\mathrm{A}}\mathbf{1}_n, & n=y_C+1,y_C+2\cdots,C.\end{cases}$$

$$\mathbf{W}_{g_{\mathrm{p53}}}^{n}=\begin{cases}\mathbf{0}, & n=0,1,\cdots,y_C-1;\\ \sum_{m=0}^{n-1}g_{\mathrm{p53}}^{(m)}P_{B\geq n-m}\mathbf{1}_{m+1}, & n=y_C,y_C+1\cdots,C.\end{cases}$$

$$\mathbf{W}_{g_{\mathrm{A}}}^{n}=\begin{cases}\sum_{m=y_C}^{C-1}g_{\mathrm{A}}\mathbf{1}_{m+1}, & n=y_C-1;\\ \mathbf{0}, & n\neq y_C-1.\end{cases}$$

Thus, the column vector $\mathbf{W}$ can be defined as the following form

$$\mathbf{W}=\mathbf{W}_{g_{\mathrm{p53}}}+\mathbf{W}_{g_{\mathrm{A}}}+\mathbf{W}_{d_{\mathrm{A}}}=\left[\mathbf{W}_0^{\mathrm{T}},\mathbf{W}_1^{\mathrm{T}},\mathbf{W}_2^{\mathrm{T}},\cdots,\mathbf{W}_n^{\mathrm{T}}\right]^{\mathrm{T}} \tag{S49}$$

where $\mathbf{W}_n=\mathbf{W}_{g_{\mathrm{p53}}}^{n}+\mathbf{W}_{g_{\mathrm{A}}}^{n}+\mathbf{W}_{d_{\mathrm{A}}}^{n}$, which can be expressed as

$$\mathbf{W}_n=\begin{cases}\mathbf{0}, & n=0,1,\cdots,y_C-2;\\ \sum_{m=y_C}^{C-1}g_{\mathrm{A}}\mathbf{1}_{m+1}, & n=y_C-1;\\ \sum_{m=0}^{n-1}g_{\mathrm{p53}}^{(m)}P_{B\geq n-m}\mathbf{1}_{m+1}, & n=y_C;\\ nd_{\mathrm{A}}\mathbf{1}_n+\sum_{m=0}^{n-1}g_{\mathrm{p53}}^{(m)}P_{B\geq n-m}\mathbf{1}_{m+1}, & n=y_C+1,y_C+2\cdots,C.\end{cases}$$

In brevity, the involved vectors can be expressed as

$$\mathbf{W}_{d_{\mathrm{A}}}^{\mathrm{T}}=\left[[0,\cdots,0],\cdots,[0,\cdots,0],\left[0,\cdots,0,\left(y_C+1\right)d_{\mathrm{A}},0,\cdots,0\right],\left[0,\cdots,0,\left(y_C+2\right)d_{\mathrm{A}},0,\cdots,0\right],\cdots,\left[0,\cdots,0,Cd_{\mathrm{A}}\right]\right]$$

$$\begin{aligned}\mathbf{W}_{g_{\mathrm{p53}}}^{\mathrm{T}}=&\left[[0,\cdots,0],\cdots,[0,\cdots,0],\left[g_{\mathrm{p53}}^{(0)}P_{B\geq y_C},g_{\mathrm{p53}}^{(1)}P_{B\geq y_C-1},\cdots,g_{\mathrm{p53}}^{(y_C-1)}P_{B\geq 1},0,\cdots,0\right],\right.\\&\left.\left[g_{\mathrm{p53}}^{(0)}P_{B\geq y_C+1},g_{\mathrm{p53}}^{(1)}P_{B\geq y_C},\cdots,g_{\mathrm{p53}}^{(y_C)}P_{B\geq 1},0,\cdots,0\right],\cdots,\left[g_{\mathrm{p53}}^{(0)}P_{B\geq C},g_{\mathrm{p53}}^{(1)}P_{B\geq C-1},\cdots,g_{\mathrm{p53}}^{(C-1)}P_{B\geq 1}\right]\right]\end{aligned}$$

$$\mathbf{W}_{g_{\mathrm{A}}}^{\mathrm{T}}=\left[[0,\cdots,0],\cdots,[0,\cdots,0],[0,\cdots,0,g_{\mathrm{A}},g_{\mathrm{A}},\cdots,g_{\mathrm{A}}],[0,\cdots,0],\cdots,[0,\cdots,0]\right]$$

$$\begin{aligned}\mathbf{W}^{\mathrm{T}}=&\left[[0,\cdots,0],\cdots,[0,\cdots,0],[0,\cdots,0,g_{\mathrm{A}},g_{\mathrm{A}},\cdots,g_{\mathrm{A}}],\left[g_{\mathrm{p53}}^{(0)}P_{B\geq y_C},g_{\mathrm{p53}}^{(1)}P_{B\geq y_C-1},\cdots,g_{\mathrm{p53}}^{(y_C-1)}P_{B\geq 1},0,\cdots,0\right],\right.\\&\left[g_{\mathrm{p53}}^{(0)}P_{B\geq y_C+1},g_{\mathrm{p53}}^{(1)}P_{B\geq y_C},\cdots,g_{\mathrm{p53}}^{(y_C-1)}P_{B\geq 2},\left(y_C+1\right)d_{\mathrm{A}}+g_{\mathrm{p53}}^{(y_C)}P_{B\geq 1},0,\cdots,0\right],\cdots,\\&\left.\left[g_{\mathrm{p53}}^{(0)}P_{B\geq C},g_{\mathrm{p53}}^{(1)}P_{B\geq C-1},\cdots,Cd_{\mathrm{A}}+g_{\mathrm{p53}}^{(C-1)}P_{B\geq 1}\right]\right]\end{aligned}$$

The formulation for calculating the moments of the FPT is the same as Eq. S17.

Numerical results for mean FPT and timing variability are shown in Fig. S7.

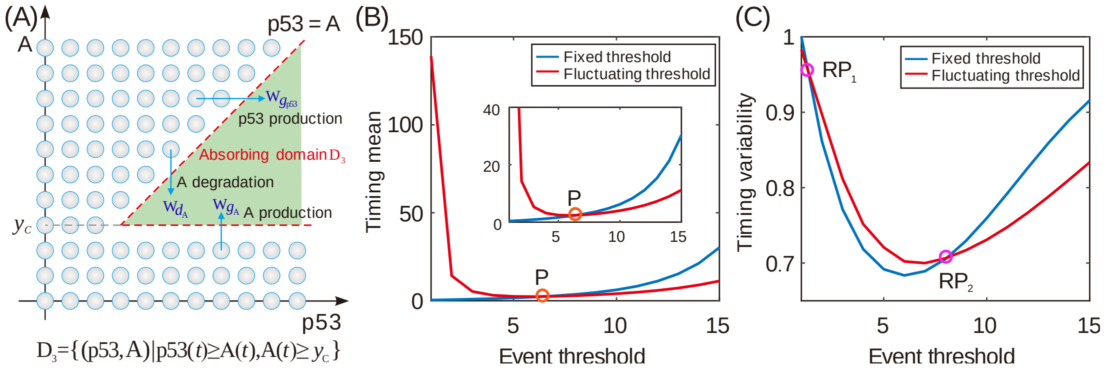

**Fig. S7 Characteristic of the curve for timing mean or timing variability vs event threshold in the case of absorbing domain** $\mathcal{D}_3$. (A) Schematic for PFT when an absorbing domain is defined by $\mathcal{D}_3 = \{(p53, A) | \, p53(t) \geq A(t), A(t) \geq y_C\}$. (B) Timing mean as a function of event threshold for two different kinds of thresholds, where there is a crossing point $\mathrm{P}$ which is the same mean with the Fig. S6*B*. (C) Timing variability as a function of event threshold for two different kinds of thresholds, where two empty circle represent the crossing point of two curves, denoted by $\mathrm{RP}_1$ and $\mathrm{RP}_2$. In (B) and (C), the parameter values are set as $g_{\mathrm{p53}} = 5$, $d_{\mathrm{p53}} = 1$, $b = 1$ and $y_{\mathrm{C}} = 5$, the cutoff constant is set as $p53_{\mathrm{max}} = 50$, and the range of threshold is set as $A_{\mathrm{thresold}} = 1 \sim 15$. If the fixed threshold is set as $A_{\mathrm{threshold}} = 10$, the fluctuating threshold corresponds to $g_{\mathrm{A}} = 10$ and $d_{\mathrm{A}} = 1$.

#### 3.4 The absorbing domain is $\mathcal{D}_4$

Here, we analyze the case of absorbing domain $\{(p53, A) | \, p53(t) \geq A(t), \, p53(t) \geq x_C, A(t) \geq y_C\}$. Obviously, it becomes the above case **S3.1** when $x_C = y_C = 0$ (referring to Fig. S3*A*), and the above case **S3.2** when $x_C > 0, y_C = 0$ (referring to Fig. S3*B*), and the above case **S3.3** if $x_C = y_C > 0$ or $y_C > x_C \geq 0$ (referring to Fig. S3*C*). However, the case of $x_C > y_C > 0$ is an exception since it does not belong to these three cases. This case will be discussed in **S3.4** of this section wherein the absorbing domain is specified as

$$\mathcal{D}_4 = \{(p53, A) | \, p53(t) \geq A(t), \, p53(t) \geq x_C, A(t) \geq y_C\} \text{ with } x_C > y_C > 0.$$

If the trajectory $(p53, A)$ of protein pair $p53$ and $A$ is considered in three domains: $\{(p53, A) | \, A(t) < y_C\}$, $\{(p53, A) | \, p53(t) < x_C, y_C \leq A(t) < x_C\}$ and $\{(p53, A) | \, p53(t) < A(t), A(t) \geq x_C\}$ (referring to Fig. S3*D*), we define the random variable $T$ by the time at which protein counts $p53$ reach protein counts $A$ for the first time, that is,

$$T = \min\left\{t : p53(t) \geq A(t), p53(t) \geq x_C, A(t) \geq y_C \middle| p53(0) = m < A(0) = n\right\} \tag{S50}$$

The absorbing domain in this case is given by

$$\mathcal{D}_4 = \left\{(p53, A) \middle| p53(t) \geq A(t),\ p53(t) \geq x_C, A(t) \geq y_C\right\} \tag{S51}$$

And the corresponding finite state-space for birth-death process $\Omega$ is given by $\Omega = \Omega_1 \cup \Omega_2 \cup \Omega_3$, where

$\Omega_1 = \left\{(p53, A) \middle| A(t) < y_C, p53(t) = 0,1,2,\cdots,C-1, A(t) = 0,1,2,\cdots,y_C - 1\right\}$,

$\Omega_2 = \left\{(p53, A) \middle| p53(t) < x_C, y_C \leq A(t) < x_C, p53(t) = 0,1,\cdots,x_C - 1, A(t) = y_C, y_C + 1,\cdots,x_C - 1\right\}$,

$\Omega_3 = \left\{(p53, A) \middle| p53(t) < A(t), A(t) \geq x_C, p53(t) = 0,1,2,\cdots,C-1, A(t) = x_C, x_C + 1, x_C + 2,\cdots,C\right\}$.

Then we easily write the finite state-space $\Omega$, which consists of three parts , i.e., $\Omega = \Omega_1 \cup \Omega_2 \cup \Omega_3$, where we denote respectively

$$\Omega_1 = \left\{(m,n) \middle| m = 0,1,2,\cdots,C-1, n = 0,1,2,\cdots,y_C - 1\right\}, \tag{S52}$$

$$\Omega_2 = \left\{(m,n) \middle| m = 0,1,\cdots,x_C - 1, n = y_C, y_C + 1,\cdots,x_C - 1\right\}, \tag{S53}$$

$$\Omega_3 = \left\{(m,n) \middle| m < n, m = 0,1,2,\cdots,C-1, n = x_C, x_C + 1, x_C + 2,\cdots,C\right\}. \tag{S54}$$

Moreover, the corresponding chemical master equation for $P_{m,n}(t)$ takes the form of Eq. S25 with $(m,n) \in \Omega$ , where we denote

$$\begin{aligned}
\mathbf{P}_{m,n}(t) &= \left[\mathbf{P}_{*,0},\cdots,\mathbf{P}_{*,y_C-1},\mathbf{P}_{*,y_C},\mathbf{P}_{*,y_C+1},\cdots,\mathbf{P}_{*,x_C},\mathbf{P}_{*,x_C+1},\cdots,\mathbf{P}_{*,C}\right]^{\mathrm{T}} \\
&= \Big[\left[P_{0,0},P_{1,0},\cdots,P_{C-1,0}\right],\cdots,\left[P_{0,y_C-1},P_{1,y_C-1},\cdots,P_{C-1,y_C-1}\right],\left[P_{0,y_C},P_{1,y_C},\cdots,P_{x_C-1,y_C},0,\cdots,0\right], \\
&\quad \left[P_{0,y_C+1},P_{1,y_C+1},\cdots,P_{x_C-1,y_C+1},0,\cdots,0\right],\cdots,\left[P_{0,x_C},P_{1,x_C},\cdots,P_{x_C-1,x_C},0,\cdots,0\right], \\
&\quad \left[P_{0,x_C+1},P_{1,x_C+1},\cdots,P_{x_C,x_C+1},0,\cdots,0\right],\cdots,\left[P_{0,\mathrm{C}},P_{1,\mathrm{C}},\cdots,P_{C-1,\mathrm{C}}\right]\Big]^{\mathrm{T}}
\end{aligned}$$

Note that $x_C > y_C$ and $\mathbf{P}_{*,k}$ has three forms,

$$\begin{cases}
\mathbf{P}_{*,\mathrm{k}} = \left[P_{0,k},P_{1,k},\cdots,P_{C-1,k}\right], & k = 0,1,\cdots,y_C - 1; \\
\mathbf{P}_{*,\mathrm{k}} = \left[P_{0,k},P_{1,k},\cdots,P_{x_C-1,k},0,\cdots,0\right], & k = y_C, y_C + 1,\cdots,x_C - 1. \\
\mathbf{P}_{*,\mathrm{k}} = \left[P_{0,k},P_{1,k},\cdots,P_{k-1,k},0,\cdots,0\right], & k = x_C, x_C + 1,\cdots,C.
\end{cases}$$

Similarity, we easily determine the matrix $\mathbf{M}$ in equation $\partial_t \mathbf{P}_{m,n}(t) = \mathbf{M}\mathbf{P}_{m,n}(t)$ with $(m,n) \in \Omega$ for the sub-matrix operator $\mathbb{L}$, that is

$$\mathbf{M}=\begin{bmatrix}\mathbf{D}_0 & \mathbf{U}_0 & & & & \\ \mathbf{L}_0 & \mathbf{D}_1 & \mathbf{U}_1 & & & \\ & \mathbf{L}_1 & \mathbf{D}_2 & \mathbf{U}_2 & & \\ & & \ddots & \ddots & \ddots & \\ & & & \mathbf{L}_{C-2} & \mathbf{D}_{C-1} & \mathbf{U}_{C-1} \\ & & & & \mathbf{L}_{C-1} & \mathbf{D}_C\end{bmatrix} \tag{S55}$$

where

$$\mathbf{U}_i=\begin{cases}(i+1)d_\mathrm{A}\mathbf{I}_C, & i=0,1,\cdots,y_C-2;\\ (i+1)\left(\mathbb{L}^{(x_C)}\left(d_\mathrm{A}\mathbf{I}_C\right)\right), & i=y_C-1,y_C,\cdots,x_C;\\ (i+1)\left(\mathbb{L}^{(i)}\left(d_\mathrm{A}\mathbf{I}_C\right)\right), & i=x_C+1,x_C+2,\cdots,C-1.\end{cases}$$

$$\mathbf{D}_i=\begin{cases}\mathbf{M}_{g_\mathrm{p53}}+\mathbf{M}_{d_\mathrm{p53}}-\left(g_\mathrm{A}+id_\mathrm{A}\right)\mathbf{I}_C, & i=0,1,\cdots,y_C-1;\\ \mathbb{L}^{(x_C)}\left(\mathbf{M}_{g_\mathrm{p53}}+\mathbf{M}_{d_\mathrm{p53}}-\left(g_\mathrm{A}+id_\mathrm{A}\right)\mathbf{I}_C\right), & i=y_C,y_C+1,\cdots,x_C;\\ \mathbb{L}^{(i)}\left(\mathbf{M}_{g_\mathrm{p53}}+\mathbf{M}_{d_\mathrm{p53}}-\left(g_\mathrm{A}+id_\mathrm{A}\right)\mathbf{I}_C\right), & i=x_C+1,x_C+2,\cdots,C.\end{cases}$$

$$\mathbf{L}_i\equiv\begin{cases}g_\mathrm{A}\mathbf{I}_C, & i=0,1,\cdots,y_C-2;\\ \mathbb{L}^{(x_C)}\left(g_\mathrm{A}\mathbf{I}_C\right), & i=y_C-1,y_C,\cdots,x_C;\\ \mathbb{L}^{(i)}\left(g_\mathrm{A}\mathbf{I}_C\right), & i=x_C+1,x_C+2,\cdots,C-1.\end{cases}$$

Here, $\mathbf{I}_C$, $\mathbf{M}_{d_\mathrm{p53}}$ and $\mathbf{M}_{g_\mathrm{p53}}$ have same definition with above case.

In this case, for the given numerical cutoff $(C)$, we can determine the probability that a state reaches the absorbing domain defined by $\mathcal{D}_4=\left\{(p53,A)\middle|\, p53(t)\ge A(t),\ p53(t)\ge x_C, A(t)\ge y_C\right\}$ with $x_C>y_C$ in the infinitesimal time interval $(t,t+dt)$ . This probability is the sum of the following four terms: the probability that $\left\{p53(t)=m, A(t)=n\right\}$ and a jump of size $n-m$ or large occurs in the time interval $[t,t+dt)$. the probability that $\left\{p53(t)=m, A(t)=n\right\}$ and a jump of size $x_C-m$ or large occurs in the time interval $[t,t+dt)$ , the degradation probability that $\left\{p53(t)=n-1, A(t)=n\right\}$ with $n\ge x_C+1$ occurs in the time interval $[t,t+dt)$, and the transcription probability that $\left\{p53(t)=m, A(t)=y_C-1\right\}$ with $m\ge x_C$ occurs in the time interval $[t,t+dt)$. Thus, the probability density function of the FPT that $(p53,A)$ reach absorbing domain $\mathcal{D}_4$ is given by

$$f_T(t) = \sum_{\substack{n=x_C+1 \\ m=n-1}}^{C} n d_A P_{m,n}(t) + \sum_{n=x_C}^{C}\sum_{m=0}^{n-1} g_{p53}^{(m)} P_{B\geq n-m} P_{m,n}(t)$$
$$+\sum_{n=y_C}^{x_C-1}\sum_{m=0}^{x_C-1} g_{p53}^{(m)} P_{B\geq x_C-m} P_{m,n}(t) + \sum_{\substack{m=x_C \\ n=y_C-1}}^{C-1} g_A P_{m,n}(t) \quad \text{(S56)}$$
$$= \mathbf{W}_{d_A}^{T}\mathbf{P}_{m,n}(t) + \mathbf{W}_{g_{p53}}^{(1)T}\mathbf{P}_{m,n}(t) + \mathbf{W}_{g_{p53}}^{(2)T}\mathbf{P}_{m,n}(t) + \mathbf{W}_{g_A}^{T}\mathbf{P}_{m,n}(t)$$
$$\equiv \mathbf{W}^{T}\mathbf{P}_{m,n}(t)$$

where we denote respectively three column vectors

$$\begin{cases} \mathbf{W}_{d_A} = \left(\mathbf{W}_{d_A}^{n}\right)_{(C+1)\times 1} \\ \mathbf{W}_{g_{p53}}^{(1)} = \left(\mathbf{W}_{g_{p53}}^{n}\right)_{(C+1)\times 1} \\ \mathbf{W}_{g_{p53}}^{(2)} = \left(\hat{\mathbf{W}}_{g_{p53}}^{n}\right)_{(C+1)\times 1} \\ \mathbf{W}_{g_A} = \left(\mathbf{W}_{g_A}^{n}\right)_{(C+1)\times 1} \end{cases}$$

with

$$\mathbf{W}_{d_A}^{n} = \begin{cases} \mathbf{0}, & n = 0,1,\cdots,x_C; \\ n d_A \mathbf{1}_n, & n = x_C+1, x_C+2\cdots,C. \end{cases}$$

$$\mathbf{W}_{g_{p53}}^{n} = \begin{cases} \mathbf{0}, & n = 0,1,\cdots,x_C-1; \\ \sum_{m=0}^{n-1} g_{p53}^{(m)} P_{B\geq n-m}\mathbf{1}_{m+1}, & n = x_C, x_C+1,\cdots,C. \end{cases}$$

$$\hat{\mathbf{W}}_{g_{p53}}^{n} = \begin{cases} \mathbf{0}, & n = 0,1,\cdots,y_C-1, x_C, x_C+1,\cdots,C; \\ \sum_{m=0}^{x_C-1} g_{p53}^{(m)} P_{B\geq x_C-m}\mathbf{1}_{m+1}, & n = y_C, y_C+1,\cdots,x_C-1. \end{cases}$$

$$\mathbf{W}_{g_A}^{n} = \begin{cases} \sum_{m=y_C}^{C-1} g_A \mathbf{1}_{m+1}, & n = y_C-1; \\ \mathbf{0}, & n \neq y_C-1. \end{cases}$$

Thus the column vector $\mathbf{W}$ can be expressed as the follow form

$$\mathbf{W} = \mathbf{W}_{g_{p53}}^{(1)} + \mathbf{W}_{g_{p53}}^{(2)} + \mathbf{W}_{g_A} + \mathbf{W}_{d_A} = \left[\mathbf{W}_0^T, \mathbf{W}_1^T, \mathbf{W}_2^T, \cdots, \mathbf{W}_n^T\right]^T \quad \text{(S57)}$$

where $\mathbf{W}_n = \mathbf{W}_{g_{p53}}^{n} + \hat{\mathbf{W}}_{g_{p53}}^{n} + \mathbf{W}_{g_A}^{n} + \mathbf{W}_{d_A}^{n}$, which can be expressed as

$$\mathbf{W}_n = \begin{cases} \mathbf{0}, & n = 0,1,\cdots,y_C-2; \\ \sum_{m=x_C}^{C-1} g_A \mathbf{1}_{m+1}, & n = y_C-1; \\ \sum_{m=0}^{x_C-1} g_{p53}^{(m)} P_{B\geq x_C-m}\mathbf{1}_{m+1}, & n = y_C, y_C+1,\cdots,x_C-1, x_C; \\ n d_A \mathbf{1}_n + \sum_{m=0}^{n-1} g_{p53}^{(m)} P_{B\geq n-m}\mathbf{1}_{m+1}, & n = x_C+1, x_C+2\cdots,C. \end{cases}$$

In brevity, the involved vectors can be expressed as

$$\mathbf{W}_{d_A}^{T} = \left[[0,\cdots,0],\cdots,[0,\cdots,0],[0,\cdots,0,(x_C+1)d_A,0,\cdots,0],\cdots,[0,\cdots,0,C d_A]\right]$$

$$\mathbf{W}_{g_x}^{(1)\mathrm{T}} = \Big[[0,\cdots,0],\cdots,[0,\cdots,0],\Big[g_{\mathrm{p53}}^{(0)}P_{B\geq x_C}, g_{\mathrm{p53}}^{(1)}P_{B\geq x_C-1},\cdots,g_{\mathrm{p53}}^{(x_C-1)}P_{B\geq 1},0,\cdots,0\Big],$$
$$\Big[g_{\mathrm{p53}}^{(0)}P_{B\geq x_C+1}, g_{\mathrm{p53}}^{(1)}P_{B\geq x_C},\cdots,g_{\mathrm{p53}}^{(x_C)}P_{B\geq 1},0,\cdots,0\Big],\cdots,\Big[g_{\mathrm{p53}}^{(0)}P_{B\geq C}, g_{\mathrm{p53}}^{(1)}P_{B\geq C-1},\cdots,g_{\mathrm{p53}}^{(C-1)}P_{B\geq 1}\Big]\Big]$$
$$\mathbf{W}_{g_x}^{(2)\mathrm{T}} = \Big[[0,\cdots,0],\cdots,[0,\cdots,0],\Big[g_{\mathrm{p53}}^{(0)}P_{B\geq x_C}, g_{\mathrm{p53}}^{(1)}P_{B\geq x_C-1}\cdots,g_{\mathrm{p53}}^{(x_C-1)}P_{B\geq 1},0,\cdots,0\Big],\cdots,$$
$$\Big[g_{\mathrm{p53}}^{(0)}P_{B\geq x_C}, g_{\mathrm{p53}}^{(1)}P_{B\geq x_C-1}\cdots,g_{\mathrm{p53}}^{(x_C-1)}P_{B\geq 1},0,\cdots,0\Big],[0,\cdots,0],\cdots,[0,\cdots,0]\Big]$$

$$\mathbf{W}_{g_y}^{\mathrm{T}} = \big[[0,\cdots,0],\cdots,[0,\cdots,0],[0,\cdots,0,\mathrm{g_A},\mathrm{g_A},\cdots,\mathrm{g_A}],[0,\cdots,0],\cdots,[0,\cdots,0]\big]$$

$$\mathbf{W}^{\mathrm{T}} = \big[[0,\cdots,0],\cdots,[0,\cdots,0],[0,\cdots,0,\mathrm{g_A},\mathrm{g_A},\cdots,\mathrm{g_A}],\Big[g_{\mathrm{p53}}^{(0)}P_{B\geq x_C}, g_{\mathrm{p53}}^{(1)}P_{B\geq x_C-1}\cdots,g_{\mathrm{p53}}^{(x_C-1)}P_{B\geq 1},0,\cdots,0\Big],\cdots,$$
$$\Big[g_{\mathrm{p53}}^{(0)}P_{B\geq x_C}, g_{\mathrm{p53}}^{(1)}P_{B\geq x_C-1}\cdots,g_{\mathrm{p53}}^{(x_C-1)}P_{B\geq 1},0,\cdots,0\Big],$$
$$\Big[g_{\mathrm{p53}}^{(0)}P_{B\geq x_C+1}, g_{\mathrm{p53}}^{(1)}P_{B\geq x_C},\cdots,g_{\mathrm{p53}}^{(x_C-1)}P_{B\geq 2},(x_C+1)d_{\mathrm{A}}+g_{\mathrm{p53}}^{(x_C)}P_{B\geq 1},0,\cdots,0\Big],\cdots,$$
$$\Big[g_{\mathrm{p53}}^{(0)}P_{B\geq C}, g_{\mathrm{p53}}^{(1)}P_{B\geq C-1},\cdots,Cd_{\mathrm{A}}+g_{\mathrm{p53}}^{(C-1)}P_{B\geq 1}\Big]\Big]$$

The formulation for calculating the moments of the FPT is the same as Eq. S17.

Numerical results for mean FPT and timing variability are shown in Fig. S8.

From Figs S6*A*, S7*A* and S8*A*, we can observe that the absorbing domains specified above are smaller than the one in Fig. S3*A*. Figures S6, S7, S8 demonstrate that fluctuations in event threshold can affect the arriving time that the regulated protein reaches a certain fluctuating threshold for the first time, and event threshold can impact the timing variability. We also observe that for a high event threshold, fluctuations in event threshold can improve the event respond and shorten the time of FPT. The corresponding variability tendency in the timing, which raises the time precision, also makes this result clear.

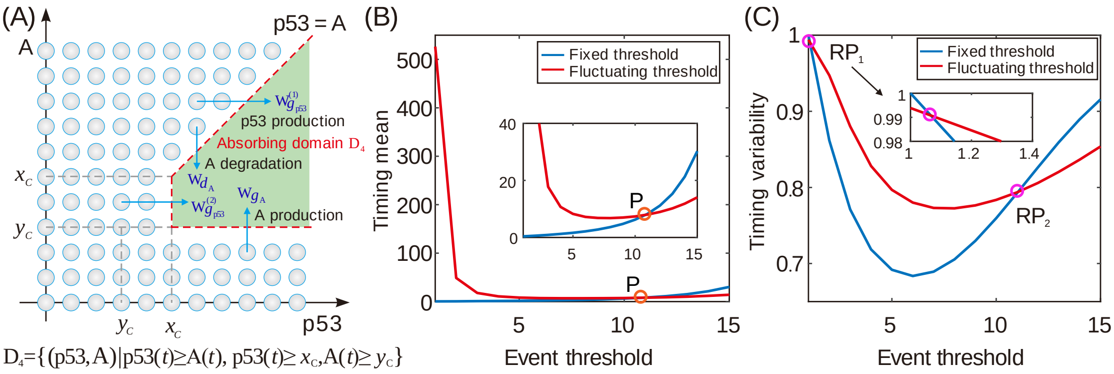

**Fig. S8 Characteristic of the curve for timing mean or timing variability vs event threshold in the case of absorbing domain** $\mathcal{D}_4$. (A) Schematic for PFT when an absorbing domain is defined by $\mathcal{D}_4 = \{(p53, A)|\, p53(t) \geq A(t),\ p53(t) \geq x_C, A(t) \geq y_C\}$ with $x_C > y_C$. (B) Timing mean as a function of event threshold for two different kinds of thresholds, where there is a crossing point $P$

which the same mean with the Fig. S6*B*. (C) Timing variability as a function of event threshold for two different kinds of thresholds, where two empty circle represent the crossing point of two curves, denoted by $\mathrm{RP}_1$ and $\mathrm{RP}_2$. In (B) and (C), the parameter values are set as $g_{\mathrm{p53}}=5$, $d_{\mathrm{p53}}=1$, $b=1$, $x_{\mathrm{C}}=10$ and $y_{\mathrm{C}}=5$, the cutoff constant is set as $p53_{\mathrm{max}}=50$, and the range of threshold is set as $A_{\mathrm{thresold}}=1\sim 15$. If the fixed threshold is set as $A_{\mathrm{threshold}}=10$, the fluctuating threshold corresponds to $g_{\mathrm{A}}=10$ and $d_{\mathrm{A}}=1$.

### 4 Effect of timescales on the timing event

In this section, we investigate the effects of timescales on timing precision and mean FPT. For simplicity, we consider the simplest case i.e., the absorbing domain is $\mathcal{D}_1$ (referring to Fig. S3*A*). We firstly define the timescale in event timing. If the production and degradation rate of protein $p53$ or $A$ are simultaneously enlarged by $\alpha_{\mathrm{p53}}$ or $\alpha_{\mathrm{A}}$ times, the factor $\alpha_{\mathrm{p53}}$ or $\alpha_{\mathrm{A}}$ is defined as the timescale of protein $p53$ or $A$. The size of factor $\alpha_{\mathrm{p53}}$ or $\alpha_{\mathrm{A}}$ usually affects fluctuations in protein $p53$ or $A$. Now, we derive analytical results on the effects of timescale factors.

Based on the definition of timescale and by matrix $\mathbf{M}$ in Eq. S31, and by simultaneously enlarging the production and degradation rate of protein $p53$ or $A$, we have

$$\begin{cases} \mathbf{U}_i=(i+1)\alpha_{\mathrm{A}}\mathbb{L}^{(i)}(d_{\mathrm{A}}\mathbf{I}_C), & i=1,2,\cdots,C-1; \\ \mathbf{D}_i=\mathbb{L}^{(i)}\left(\alpha_{\mathrm{p53}}\mathbf{M}_{g_{\mathrm{p53}}}+\alpha_{\mathrm{p53}}\mathbf{M}_{d_{\mathrm{p53}}}-\alpha_{\mathrm{A}}(g_{\mathrm{A}}+id_{\mathrm{A}})\mathbf{I}_C\right), & i=1,2,\cdots,C; \\ \mathbf{L}_i=\alpha_{\mathrm{A}}\mathbb{L}^{(i)}(g_{\mathrm{A}}\mathbf{I}_C), & i=1,2,\cdots,C-1. \end{cases}$$

Thus $\mathbf{M}$ can be rewritten as $\mathbf{M}=\alpha_{\mathrm{p53}}\mathbf{M}_1+\alpha_{\mathrm{A}}\mathbf{M}_2$, where $\mathbf{M}_1=diag\left(\mathbf{D}_1^{\mathrm{p53}},\mathbf{D}_2^{\mathrm{p53}},\cdots,\mathbf{D}_C^{\mathrm{p53}}\right)$,

$$\mathbf{M}_2=\begin{bmatrix} \mathbf{D}_1^{\mathrm{A}} & \mathbf{U}_1 & & & & \\ \mathbf{L}_1 & \mathbf{D}_2^{\mathrm{A}} & \mathbf{U}_2 & & & \\ & \mathbf{L}_2 & \mathbf{D}_3^{\mathrm{A}} & \mathbf{U}_3 & & \\ & & \ddots & \ddots & \ddots & \\ & & & \mathbf{L}_{C-2} & \mathbf{D}_{C-1}^{\mathrm{A}} & \mathbf{U}_{C-1} \\ & & & & \mathbf{L}_{C-1} & \mathbf{D}_C^{\mathrm{A}} \end{bmatrix}$$

with $\mathbf{D}_i^{\mathrm{p53}}=\mathbb{L}^{(i)}\left(\mathbf{M}_{g_{\mathrm{p53}}}+\mathbf{M}_{d_{\mathrm{p53}}}\right)$, $\mathbf{D}_i^{\mathrm{A}}=\mathbb{L}^{(i)}\left(-(g_{\mathrm{A}}+id_{\mathrm{A}})\mathbf{I}_C\right)$, $i=1,2,\cdots,C.$

The inverse matrix of $\mathbf{M}$ is then $\mathbf{M}^{-1}=\left(\alpha_{\mathrm{p53}}\mathbf{M}_1+\alpha_{\mathrm{A}}\mathbf{M}_2\right)^{-1}$. Regarding Eq. S19, we can deduce the following expressions

$$\mathrm{CV}_T = \frac{2\mathbf{e}^{\mathrm{T}}\left(\mathbf{M}^{-1}\right)^2 \mathbf{P}(0)}{\left[\mathbf{e}^{\mathrm{T}}\mathbf{M}^{-1}\mathbf{P}(0)\right]^2} - 1 = \frac{2\mathbf{e}^{\mathrm{T}}\left(\alpha_{\mathrm{p53}}\mathbf{M}_1 + \alpha_{\mathrm{A}}\mathbf{M}_2\right)^{-2} \mathbf{P}(0)}{\left[\mathbf{e}^{\mathrm{T}}\left(\alpha_{\mathrm{p53}}\mathbf{M}_1 + \alpha_{\mathrm{A}}\mathbf{M}_2\right)^{-1}\mathbf{P}(0)\right]^2} - 1$$

$$= \frac{2\mathbf{e}^{\mathrm{T}}\left(\mathbf{M}_1 + \frac{\alpha_{\mathrm{A}}}{\alpha_{\mathrm{p53}}}\mathbf{M}_2\right)^{-2} \mathbf{P}(0)}{\left[\mathbf{e}^{\mathrm{T}}\left(\mathbf{M}_1 + \frac{\alpha_{\mathrm{A}}}{\alpha_{\mathrm{p53}}}\mathbf{M}_2\right)^{-1}\mathbf{P}(0)\right]^2} - 1 = \frac{2\mathbf{e}^{\mathrm{T}}\left(\mathbf{M}_1 + \gamma\mathbf{M}_2\right)^{-2} \mathbf{P}(0)}{\left[\mathbf{e}^{\mathrm{T}}\left(\mathbf{M}_1 + \gamma\mathbf{M}_2\right)^{-1}\mathbf{P}(0)\right]^2} - 1$$

Thus, the variability in the timing is calculated according to

$$\mathrm{CV}_T = \frac{2\mathbf{e}^{\mathrm{T}}\left(\gamma\mathbf{M}_1 + \mathbf{M}_2\right)^{-2} \mathbf{P}(0)}{\left[\mathbf{e}^{\mathrm{T}}\left(\gamma\mathbf{M}_1 + \mathbf{M}_2\right)^{-1}\mathbf{P}(0)\right]^2} - 1 \tag{S59}$$

with $\gamma = \alpha_{\mathrm{A}} / \alpha_{\mathrm{p53}}$. Eq. S59 shows how timing variability depends on the rate $\gamma$ between the timescales of proteins $p53$ and $A$. This dependence implies that variability timing does not depend on the timescale of proteins $p53$ or $A$, but depends on rate $\gamma$.

#### 5 FPT statistics in the case that burst size follows Poisson distribution

The above minimal model considers that burst size $B$ follows a geometric distribution. Here, we consider that $B$ follows a Poisson distribution. We focus on how fluctuations affect mean FPT and timing variability. Number results are shown in Figs. S9 and S10. We observe that these results are analogous to those obtained above, implying that burst size distributions have little influence on mean FPT and timing variability.

For other three cases where burst size follow a Poisson distribution, the mean FPT and timing variability also have absorbing domains and change trends similar to those in the above respective three cases, as the fluctuating threshold increases. Numerical results are shown in Figs. S11, S12, and S13.

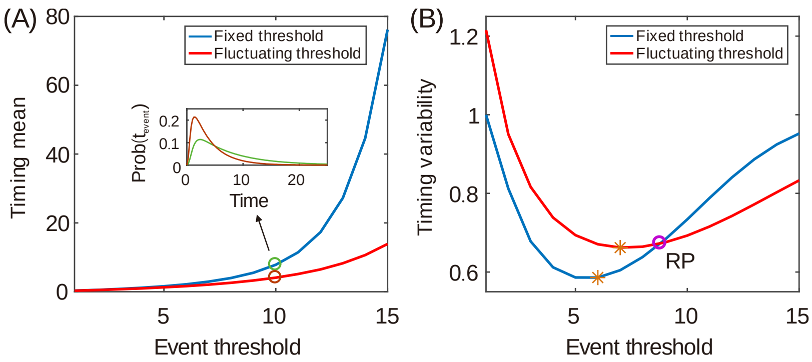

**Fig. S9 Characteristic of the curve for timing mean or timing variability vs event threshold in the case of absorbing domain** $\mathcal{D}_1$, where burst size follows a Poisson distribution. (A) Timing mean as a function of event threshold for two different kinds of thresholds, where the inset shows FPT distributions for a specific event threshold (indicated by empty circles) corresponding to $A_{\text{thresold}} = 10$. (B) Timing variability as a function of event threshold for two different kinds of thresholds, where the empty circle (denoted by RP) represents the crossing point of two curves and the responding threshold is $A_{\text{thresold}} \approx 8.8$. And stars represent the critical threshold that makes the variability in the timing reach the minimum. Variability timing for the fixed threshold is minimum when $A_{\text{thresold}} = 6$, but variability timing for the fluctuating threshold is minimum, when $A_{\text{thresold}} = 7$. In (A) and (B), the parameter values are set as $g_{\text{p53}} = 5$, $d_{\text{p53}} = 1$, $b = 1$, the cutoff constant is set as $p53_{\max} = 40$. If the fixed threshold is set as $A_{\text{threshold}} = 10$, and the fluctuating threshold corresponds to $g_{\text{A}} = 10$ and $d_{\text{A}} = 1$.

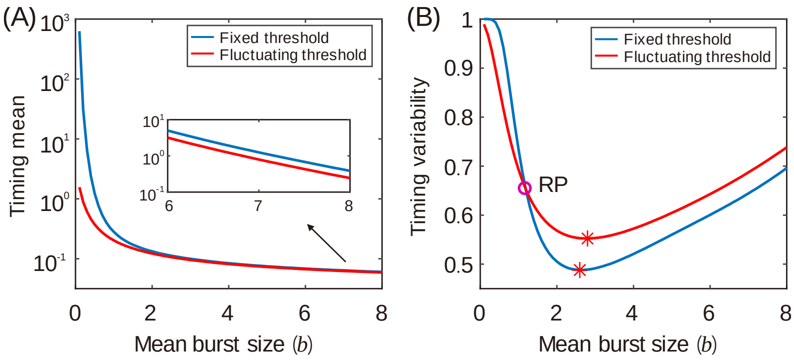

**Fig.S10 Characteristic of the curve for timing mean or timing variability vs event threshold in the case of absorbing domain** $\mathcal{D}_1$. (A) Timing mean as a function of the burst size, where the inset shows a partial enlarged diagram. (B) Timing variability as a function of the burst size, where RP represents a critical point for which the corresponding mean burst size is $b \approx 1.2$. Stars represent the critical mean burst size that makes the variability in the timing reach the minimum, which is respectively, $b \approx 2.6$ for the fixed threshold, and $b \approx 2.8$ for the fluctuating threshold. In (A) and (B), the parameter values are set as $g_{\text{p53}} = 5$, $d_{\text{p53}} = 1$, $g_{\text{A}} = 10$ and $d_{\text{A}} = 1$. The cutoff constant is set as $p53_{\max} = 40$. The range of mean burst size is set as $b = 0.1 \sim 8$, and the threshold is set as $A_{\text{thresold}} = 10$.

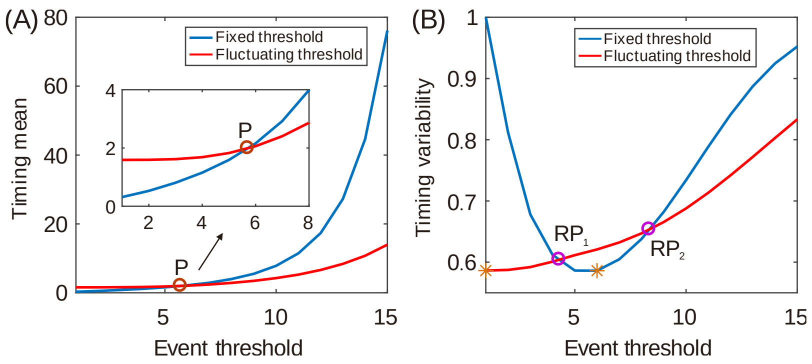

**Fig. S11 Characteristic of the curve for timing mean or timing variability vs event threshold in the case of absorbing domain** $\mathcal{D}_2$, where burst size follows a Poisson distribution. (A) Timing mean as a function of event threshold, where the inset shows a partial enlarged diagram. (B) Timing variability as a function of event threshold, where the two empty circles represent the crossing point of two curves, denoted by $\mathrm{RP}_1, \mathrm{RP}_2$. And stars represent the critical threshold that makes the variability in the timing reach the minimum, that is $A_{\mathrm{thresold}} = 6$, for the fixed threshold, but $A_{\mathrm{thresold}} = 1$, for the fluctuating threshold. Burst size follows a Poisson distribution in (A) and (B). In (A) and (B), the parameter values are set as $g_{\mathrm{p53}} = 5$, $d_{\mathrm{p53}} = 1$, $b = 1$ and $x_C = 5$, the cutoff constant is set as $p53_{\max} = 50$. If the fixed threshold is set as $A_{\mathrm{threshold}} = 10$, the fluctuating threshold corresponds to $g_{\mathrm{A}} = 10$ and $d_{\mathrm{A}} = 1$.

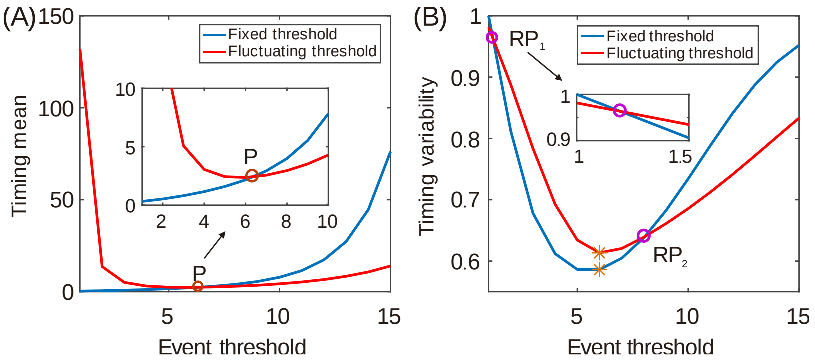

**Fig. S12 Characteristic of the curve for timing mean or timing variability vs event threshold in the case of absorbing domain** $\mathcal{D}_3$, where burst size follows Poisson distribution**.** (A) Timing mean as a function of event threshold, where the inset shows a partial enlarged diagram. (B) Timing variability as a function of event threshold, where the two empty circles represent the crossing point of two curves, denoted by $\mathrm{RP}_1, \mathrm{RP}_2$ .And stars represent the critical threshold that makes the variability in the timing reach the minimum, that is, $A_{\mathrm{thresold}} = 6$, for the fixed and fluctuating threshold. Burst size follows a Poisson distribution in (A) and (B). In (A) and (B), the parameter values are set as $g_{\mathrm{p53}} = 5$, $d_{\mathrm{p53}} = 1$, $b = 1$ and $y_C = 5$. The cutoff constant is set as $p53_{\max} = 50$. If the fixed threshold is set as $A_{\mathrm{threshold}} = 10$, the fluctuating threshold corresponds to $g_{\mathrm{A}} = 10$ and $d_{\mathrm{A}} = 1$.

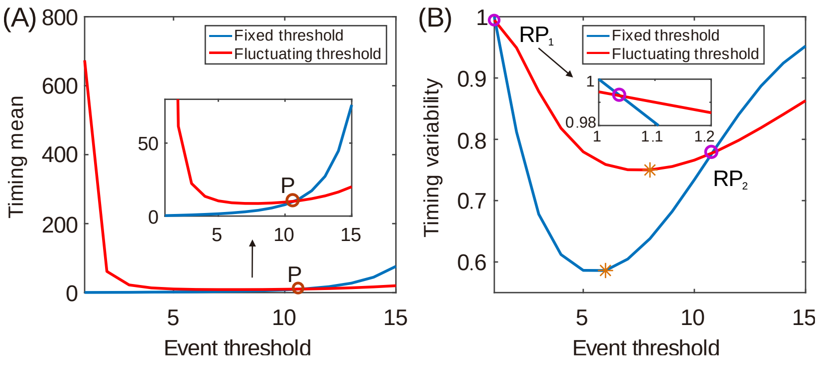

**Fig. S13 Characteristic of the curve for timing mean or timing variability vs event threshold in the case of absorbing domain** $\mathcal{D}_4$, where burst burst size follows Poisson distribution. (A) Timing mean as a function of event threshold, where the inset shows a partially enlarged diagram. (B) Timing variability as a function of event threshold, where the two empty circles represent the crossing point of two curves, denoted by $\mathrm{RP}_1, \mathrm{RP}_2$. And stars represent the critical threshold that makes the variability in the timing reach the minimum, that is, $A_{\mathrm{thresold}} = 6$ for the fixed threshold, but $A_{\mathrm{thresold}} = 8$ for the fluctuating threshold. Burst size follows a Poisson distribution in (A) and (B). In (A) and (B), parameter values are set as $g_{\mathrm{p53}} = 5$, $d_{\mathrm{p53}} = 1$, $b = 1$, $x_C = 10$, $y_C = 5$, the cutoff constant is set as $p53_{\max} = 50$. If the fixed threshold is set as $A_{\mathrm{threshold}} = 10$, the fluctuating threshold corresponds to $g_{\mathrm{A}} = 10$ and $d_{\mathrm{A}} = 1$.

[4] Van Kampen N. 2007. Stochastic process in physics and chemistry. 3rd ed. North-Holland,

Amsterdam.
